# Rescuing functional defects in a zebrafish model of CDKL5 deficiency disorder: Contribution to the identification of new therapeutic compounds

**DOI:** 10.64898/2026.03.12.711124

**Authors:** Tatiana Varela, Débora Varela, João Santos, Ana Hernández, Max Domingues, Vanessa Pinto, Natércia Conceição, M. Leonor Cancela

## Abstract

Mutations in the *CDKL5* gene cause CDKL5 deficiency disorder (CDD), a severe neurodevelopmental encephalopathy characterized by a broad range of symptoms, including early-onset seizures, profound motor impairment and dysmorphic facial features. Current treatment options remain limited and largely focus on seizure management, which is often challenging to control, underscoring the critical need for new effective therapies. To identify potential novel candidate molecules for the treatment of CDD, we performed the first *in vivo* drug screening using a *cdkl5* mutant zebrafish model. Recapitulating key features of the human disorder, *cdkl5*^-/-^ larvae exhibit reduced locomotor behavior, providing a robust readout to assess therapeutic efficacy. By screening 170 compounds from MAPK Inhibitor and Histone Modification Libraries, both implicated in CDKL5 dysfunction, we identified 18 and 12 small molecules that partially or fully restored locomotor activity, respectively. Among these, fisetin, divalproex, resveratrol, and VX-702 were further evaluated for their capacity to rescue *cdkl5*^-/-^ craniofacial defects and altered gene expression. Fisetin demonstrated the most consistent phenotypic improvement, including partial restoration of craniofacial abnormalities and normalization of gene expression levels. Future research aimed at elucidating the molecular mechanisms underlying the observed rescue effects will be critical to understand their mode of action. Overall, our study demonstrates the utility of this rapid and scalable zebrafish-based screening approach for therapeutic discovery in CDD and identifies promising therapeutic molecules that warrant further validation in complementary preclinical systems.

## 1. INTRODUCTION

Despite remarkable advances in biomedical research over recent years, approximately 95% of rare diseases still have no cure or lack effective treatment^1^. The development of new drugs suitable for therapeutic applications is particularly challenging for rare diseases due to the small population of patients, poorly understood pathophysiological mechanisms, phenotypic variability, high development costs and at limited profit potential^2–4^. Most of these conditions have a genetic origin and typically arise from mutations in single genes^1,5^.

Among rare diseases is cyclin-dependent kinase-like 5 (CDKL5) deficiency disorder (CDD), a X-linked neurodevelopmental condition caused by loss-of-function mutations in the *CDKL5* gene, which encodes a serine/threonine kinase essential for proper neuronal development and function^6–8^. CDD is a complex disorder encompassing a wide range of clinical manifestations with severity varying among affected individuals. Key features include early-onset and drug-resistant seizures, severe developmental delay, and significant motor and cognitive impairments. Other comorbidities such as hypotonia, microcephaly, dysmorphic facial features, visual impairment and scoliosis are also associated with CDD^6–10^. Currently, there is no cure for individuals affected by CDD, and available treatments are limited to alleviating symptoms, with a particular focus on seizures control, often with limited success^11–13^. Conventional antiseizure medications such as clobazam, valproate, topiramate, levetiracetam, and vigabatrin are the most frequently prescribed, however their efficacy declines over time^12,14,15^. Recently, ganaxolone became the first medication approved by both the U.S. Food and Drug Administration (FDA) and the European Medicines Agency (EMA) for the specific treatment of seizures in individuals with CDD aged 2 years and older^15–17^. Ganaxolone is a neuroactive steroid that acts as a positive allosteric modulator of GABA_A_ receptors, thus enhancing GABAergic inhibitory signaling^18,19^. Despite therapeutic advances, effective disease-modifying treatments for CDD are still lacking, reinforcing the need for pre-clinical research to identify novel therapeutic candidates capable of targeting the underlying CDD pathophysiology beyond seizure control.

Zebrafish has proven to be a robust model organism extensively used in biomedical research to model a wide range of human disorders and has become particularly valuable for high-throughput screening of small molecules^20–23^. Zebrafish produce a large number of offspring and embryos and larvae are small, enabling their use in multi-well plates, therefore facilitating the large-scale screening. Due to their translucent appearance in early developmental stages, internal organs and drug-induced phenotypic changes can be easily observed. Moreover, their capacity to absorb compounds directly from the water allows simple, cost-effective testing without invasive administration. All these advantages, in addition to a high genetic similarity with humans, make zebrafish a powerful model for preclinical drug discovery^20,23–26^.

In previous studies, our group and Serrano et al. validated a zebrafish *cdkl5* mutant line (sa21938) as a relevant preclinical model for CDKL5 deficiency disorder. This mutant showed altered motor behavior, increased susceptibility to seizures, and skeletal defects, thereby recapitulating key features of the human disorder^27,28^. Moreover, it demonstrated dysregulation of genes involved in CDKL5-associated functions in mammals^29^. Given the potential of the *cdkl5* zebrafish mutant to be used as a platform for drug discovery, in the present study we used this model to screen two libraries of small molecules, with the objective of identifying compounds capable of completely or partially reversing the mutant phenotype. Such compounds may represent promising candidates for the development of novel therapeutic strategies for CDD.

## 2. MATERIALS AND METHODS

### 2.1. Ethics statement

All procedures involving zebrafish experimentation were performed in accordance with the EU and Portuguese legislation for animal experimentation and welfare (Directives 86/609 CEE and 2010/63/EU; Decreto-Lei 113/2013; Portaria1005/92, 466/95 and 1131/97). Animal handling and experimentation were performed by qualified operators accredited by the Portuguese Direção-Geral de Alimentação e Veterinária (authorization no. 0421/2021). All efforts were made to minimize pain, distress, and discomfort.

### 2.2. Fish maintenance

Mutant *cdkl5*^sa21938^ zebrafish line generated by Stemple lab (Busch-Nentwich et al. 2013)^30^ was obtained from Zebrafish International Resource Center (ZIRC). Adult zebrafish were maintained at the CCMAR zebrafish facility in a recirculating aquatic system (Aquaneering, Inc., San Diego, CA, USA) under controlled conditions: temperature 28 ± 0.1 °C, pH 7.5 ± 0.1, conductivity 700 ± 50 μS/cm. Fish system water consisted of reverse osmosis–treated water supplemented with salt mixture (35 g/L; Instant Ocean, Blacksburg, VA, USA) and sodium bicarbonate (30 g/L; Fisher Scientific, MA, USA) to achieve the desired pH and conductivity. Zebrafish were housed under a 14 h light/10 h dark photoperiod and fed twice a day with Zebrafeed dry food (Sparos Lda, Olhão, Portugal).

For reproduction, adult females were separated from males overnight using plastic dividers placed within breeding tanks. Spawning was induced the following morning by removing the dividers, allowing natural mating to occur. The embryos were collected and raised in system water with 0.1% methylene blue to avoid fungal contamination and maintained in an incubator at 28 °C under the same photoperiod conditions as the zebrafish facility. Developmental stages were determined based on the morphological classification described by Kimmel et al. (1995)^31^. Dead or undeveloped embryos were removed daily. For the experiments, homozygous *cdkl5*^sa21938^ mutants (*cdkl5*^-/-^) and wild-type (WT) siblings’ larvae were obtained by incross of adult’s homozygous mutants and WT siblings, respectively.

### 2.3. Small molecule libraries

Two compound libraries were used in this study: the MAPK Inhibitor Library (258 compounds, catalog no. L3400) and the Histone Modification Library (347 compounds, catalog no. L4900), both purchased from Selleckchem (Houston, USA). Compounds were supplied in microplates at a stock concentration of 10 mM in dimethyl sulfoxide (DMSO) and stored at −80 °C until use.

### 2.4. Drug treatment

Solifenacin, crenigacestat, ivabradine, AZD1080, and ganaxolone were prepared as 25 and 10 mM stock solutions in DMSO and further diluted in system water to final concentration of 25 µM and 10 µM. For libraries screening assays, compound stock solutions were diluted in system water to final concentration of 10 µM, resulting in a final DMSO concentration of 0.1%, which does not affect zebrafish embryos/larvae. At 3 days post-fertilization (dpf), *cdkl5*^-/-^ larvae were transferred to 6-well plates (24 larvae per well) and exposed to compounds for 48 hours. For each experiment, WT siblings and *cdkl5*^-/-^ larvae treated with 0.1% DMSO were used as vehicle controls. Following treatment, larvae were collected for locomotor behavior analysis.

Selected hit candidate compounds-fisetin, divalproex, resveratrol, and VX-702- were also additionally administered to WT siblings to assess whether the rescue effect on the locomotor behavior was specific to the *cdkl5* mutant phenotype. These compounds were purchased individually and prepared as 10 mM stock solutions in DMSO, to confirm the positive effects observed in behavioral experiments (data not shown) and to further evaluate their ability to rescue craniofacial and molecular alterations in *cdkl5*^-/-^ larvae. Resveratrol was order from MedChemExpress (HY-16561/CS-1050), fisetin from Biorbyt (orb593928), divalproex sodium from MyBioSource (MBS579016) and VX-702 from MedChemExpress (HY-10401/CS-0074).

### 2.5. Locomotor behavioral analysis

After treatment, spontaneous swimming activity was assessed in 5 dpf fish. Larvae were randomly distributed into 24-well plates, with each well containing a single fish in 2 ml of system water. Larvae were maintained under uniform illumination, and their locomotor activity was recorded for 5 min using the Zantiks MWP automated tracking system (Zantiks Ltd., Cambridge, UK). Total distance traveled was quantified, and larvae that did not exhibit swimming behavior were excluded from the analysis. The swimming behavior of treated *cdkl5*^-/-^ larvae was compared with that of DMSO-treated wild-type (WT) and *cdkl5*^-/-^ control groups.

### 2.6. Alcian blue staining and morphometric analysis

Larvae treated with different compounds were collected and euthanized using a lethal dose of tricaine methanesulfonate (MS-222) anesthesia. Following two washes with phosphate-buffered saline (PBS), larvae were fixed overnight in 4% paraformaldehyde (PFA; Sigma). Fixed samples were washed twice with PBS and dehydrated through a graded ethanol series. Acid-free cartilage staining was performed according to the protocol of Walker and Kimmel^32^. Briefly, larvae were incubated overnight at room temperature under agitation in staining solution (0.02% Alcian blue, 40 mM MgCl₂). The pigmentation was removed by exposing the larvae to a bleaching solution of 1.5% H_2_O_2_ and 1% KOH for 20 min. Then, the tissue was cleared by sequential incubations in increasing concentrations of glycerol in KOH. Stained larvae were visualized and imaged using a stereomicroscope (Leica MZ10F) and craniofacial parameters, including ceratohyal cartilage length (Ch-l), palatoquadrate length (Pq-l), and ceratohyal cartilage width (Ch-w), were measured using Image J software.

### 2.7. RNA extraction and cDNA synthesis

Pools of *cdkl5*^-/-^ (treated with small molecules or DMSO) and WT sibling larvae (treated with DMSO) at 5 dpf were collected, euthanized with a lethal dose of MS-222 anesthesia, and washed twice with PBS. Total RNA was extracted using NZYol (NZYTech) according to the manufacturer’s instructions, and RNA quality was assessed by agarose gel electrophoresis stained with GreenSafe (NZYTech). Each experimental group consisted of three biological replicates, each prepared from pools of 24 larvae. For reverse transcription, 1 μg of total RNA was treated with RQ1 RNase-free DNase (Promega) for 30 min at 37 °C, and cDNA was synthesized by the Moloney murine leukemia virus reverse transcriptase (Invitrogen) using an oligo-dT primer.

### 2.8. Quantitative real-time PCR (qPCR)

The qPCR was performed in 20 μL reactions containing 1× NZYSpeedy qPCR Green Master Mix ROX plus (NZYTech), 0.4 μM of each forward and reverse specific primer, and 2 μL of cDNA (1:10 dilution). Primer sequences are listed in Table 1. The qPCR assays were conducted on a CFX Connect Real-Time PCR Thermocycler (Bio-Rad) under the following cycling conditions: an initial denaturation at 95°C for 2 min, followed by 40 cycles of 95°C for 5 s and 60°C for 20 s. Relative gene expression levels were calculated using the 2^−ΔΔCt^ method, with β-actin as the reference gene (Table 1).

**Table 1.**
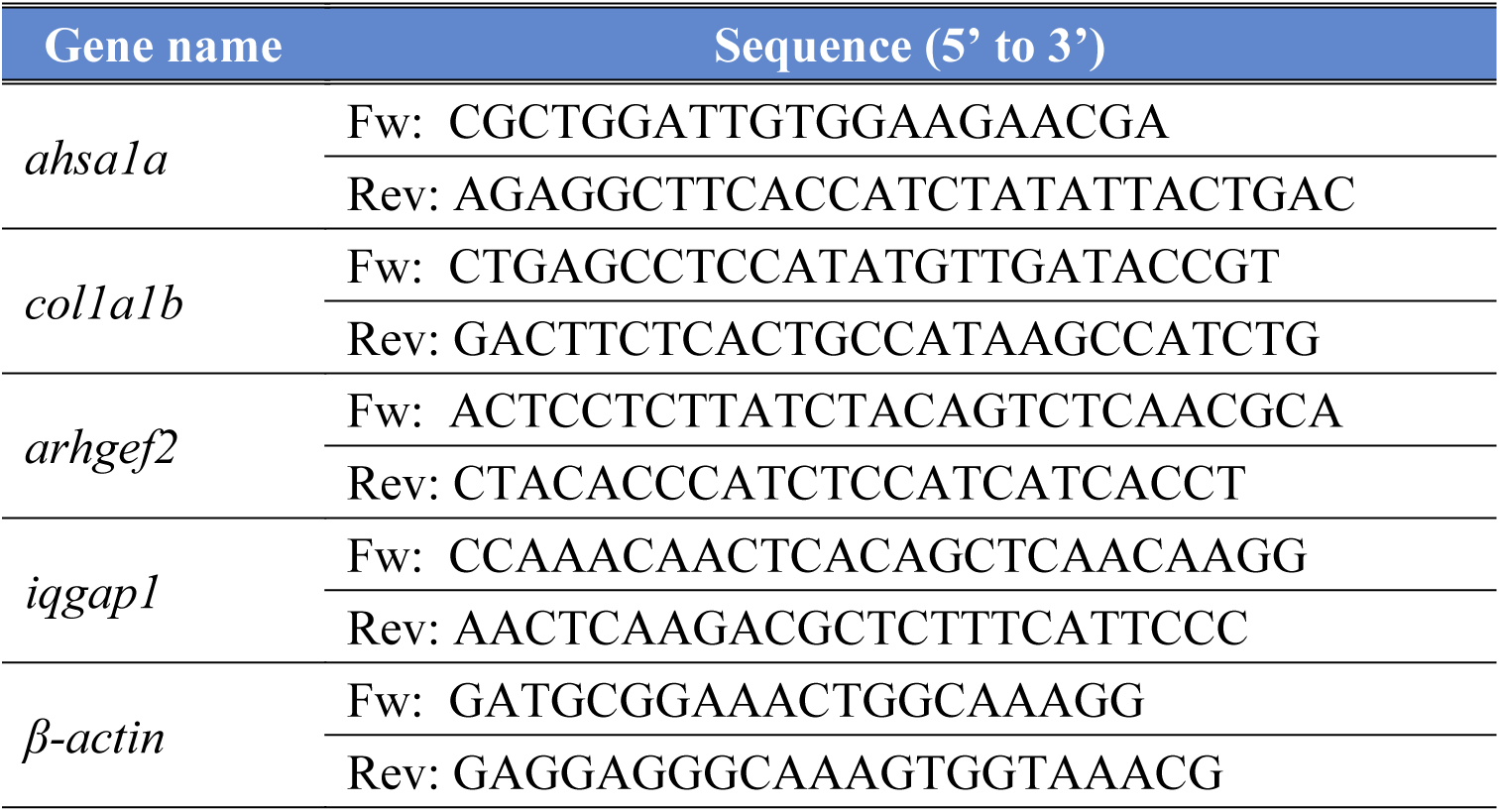
List of the primers used in this work. Fw: forward primer; Rev: reverse primer.

### 2.9. Statistical analysis

Statistical analysis was performed using Prism 5 (GraphPad software). Data normality was assessed using Kolmogorov-Smirnov test. Normally distributed data are presented as mean±SEM and significant differences were determined using one-way analysis of variance (ANOVA) for comparisons among more than two groups. Data failing the normality test are presented as median with the interquartile region (boxplot) and significant differences were determined through Kruskal-Wallis followed by Dunn multiple comparisons. Differences were considered statistically significant for p<0.05.

## 3. RESULTS

### 3.1. *cdkl5*^-/-^ mutant zebrafish as drug-screening model

To discover novel therapeutic compounds for CDKL5 deficiency disorder, we conducted a phenotype-based screening using the *cdkl5*^-/-^ mutant zebrafish. Behavioral analysis using automated locomotion tracking was used as a primary readout to identify compounds capable of restoring the abnormal swimming behavior of *cdkl5^-/-^* larvae. For this purpose, *cdkl5^-/-^* embryos at 3 dpf were subjected to treatment with the candidate compounds for two days, and their total distance traveled was analyzed and compared with that of controls. Compounds that induced a statistically significant increase in the distance traveled by *cdkl5*^-/-^ larvae compared with DMSO-treated *cdkl5*^-/-^ larvae and restored locomotor activity to levels not significantly different from WT were classified as great candidates. Compounds that improved mutant larvae locomotion but remained significantly different from wild-type (WT) larvae were classified as good candidates. A schematic illustration of the experimental screening workflow is shown in Figure 1.

**Figure 1.**
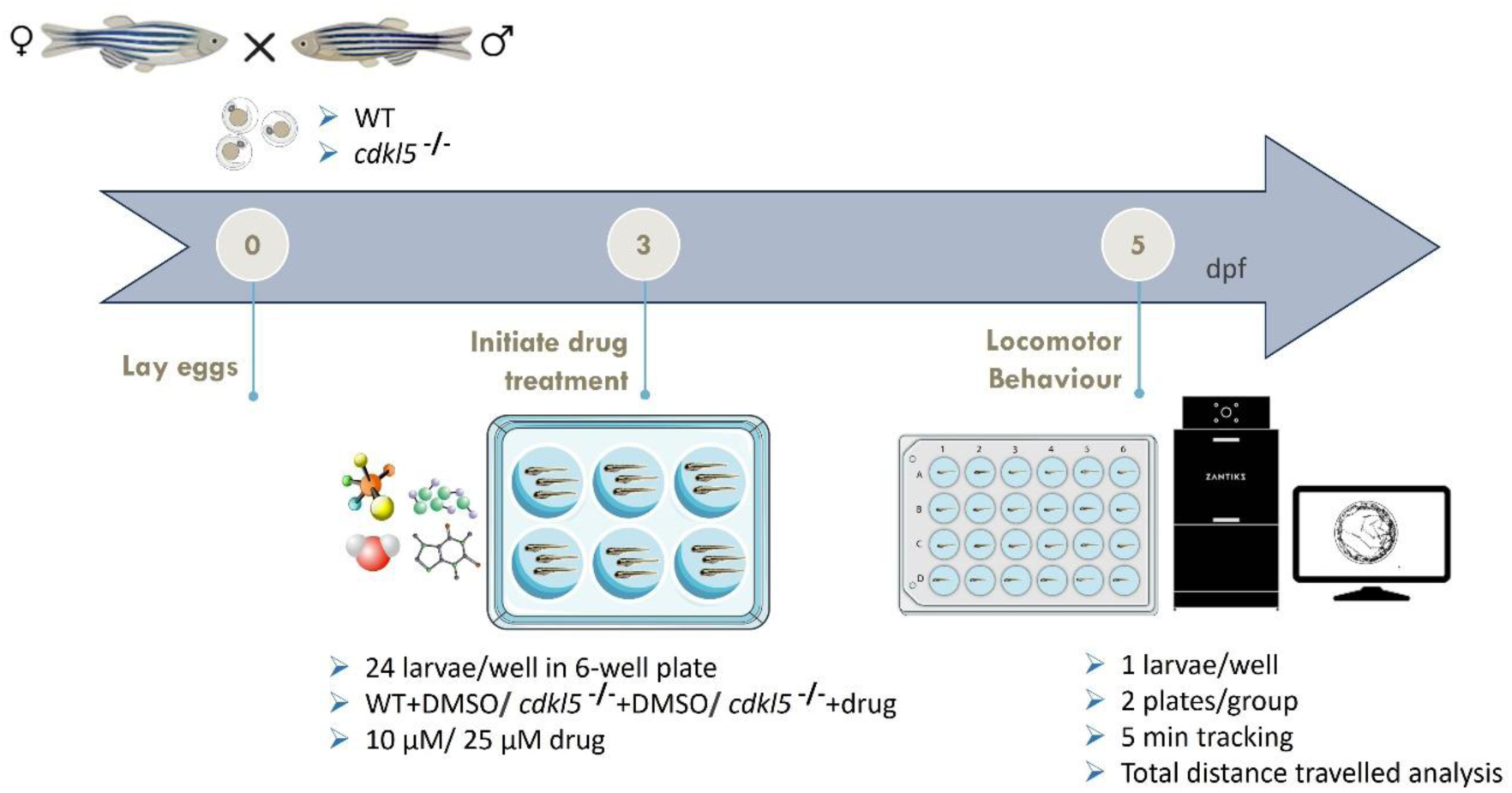
Schematic representation of the screening platform used to identify compounds capable of rescuing locomotor behavior in *cdkl5*^-/-^ zebrafish larvae.

As an initial validation of the screening approach, we first tested five compounds previously reported to rescue CDD-associated phenotypes: solifenacin, crenigacestat, ivabradine, AZD1080, and ganaxolone^16,33^. Each compound was evaluated at two concentrations (10 and 25 µM) to assess concentration-dependent effects (Figure 2). As expected, DMSO-treated *cdkl5*^⁻/⁻^ larvae exhibited a significant reduction in distance traveled compared to DMSO-treated WT. Most of the molecules at concentration of 25 µM had a negative effect on locomotor activity of *cdkl5*^⁻/⁻^ larvae. Administration of solifenacin at 10 µM induced the complete rescue of *cdkl5*^⁻/⁻^ larvae locomotor activity to WT levels. Similar, ivabradine (10 and 25 µM) and crenigacestat (10 µM) significantly increased the distance traveled by *cdkl5*^⁻/⁻^ larvae although still significantly different from WT fish. In contrast, ganaxolone and AZD1080 did not significantly improve locomotor behavior in this assay (Figure 2).

**Figure 2.**
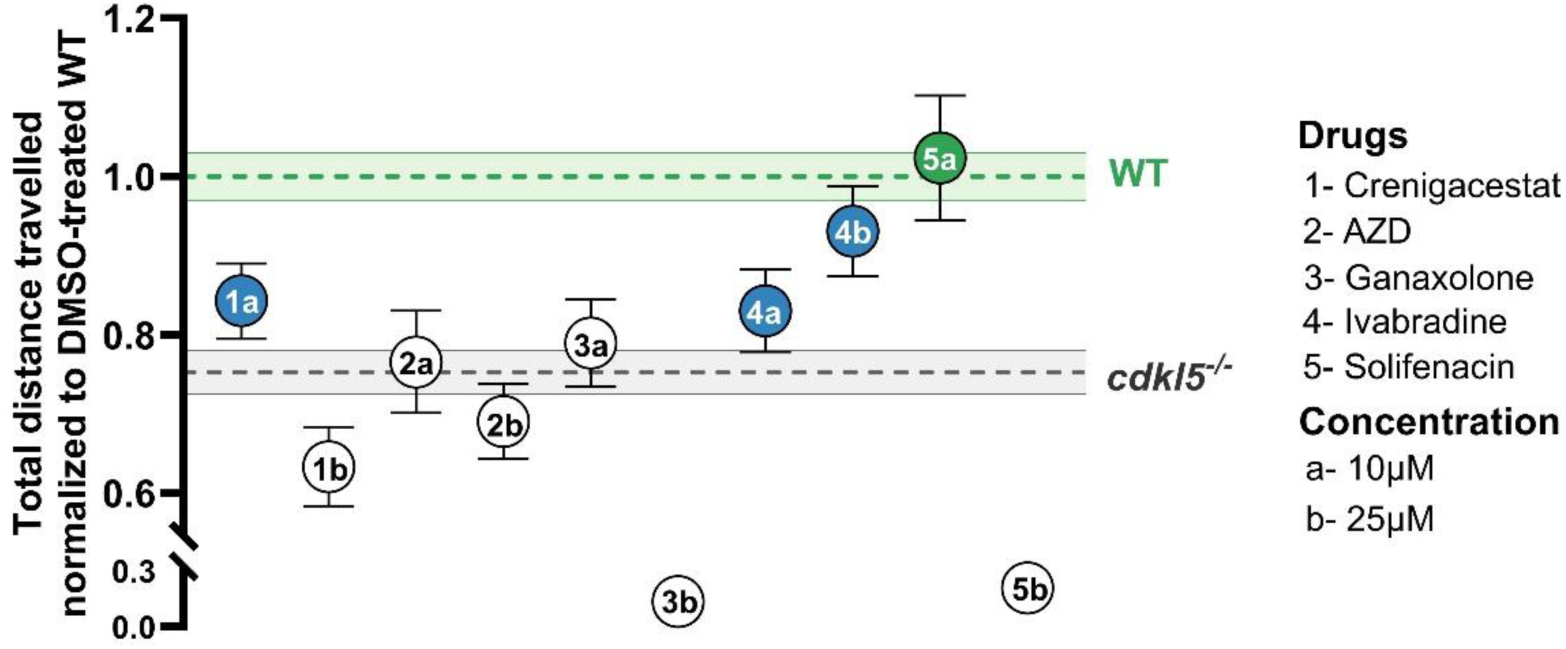
Assessment of putative CDD candidate compounds for rescue of swimming behavior in *cdkl5*^-/-^ larvae. The total distance traveled by 5 days-post fertilization (dpf) *cdkl5^-/-^* larvae treated with 10 μM or 25 μM of the candidate compounds was analyzed and normalized to wild-type (WT)*-* larvae treated with 0.1% DMSO. Each dot represents the mean ± SEM of 48 larvae. Statistical analysis was performed using one-way ANOVA followed by Dunnett’s post-hoc test. Green dots indicate compounds that differ significantly *(p*<0.05) from DMSO-treated *cdkl5^-/-^* but not from DMSO-treated WT controls (*p*>0.05). Blue dots indicate compounds that differ significantly (*p*<0.05) from both DMSO-treated *cdkl5^-/-^*and DMSO-treated WT groups.

Overall, these results support the validity of our screening approach for CDD phenotypic drug discovery.

### 3.2. Screening of two small molecule libraries to rescue *cdkl5^-/-^* locomotion behavior

To identify small molecules capable of rescuing the locomotor deficits observed in the *cdkl5*^-/-^ zebrafish model, we screened two commercially available compound libraries using our behavioral screening approach: the MAPK Inhibitor Library (L3400, Selleckchem) and the Histone Modification Library (L4900, Selleckchem). In total, 170 compounds (Supplementary Table S1) were evaluated for their ability to significantly increase the total distance traveled by 5 dpf *cdkl5*^-/-^ larvae to levels comparable to those of WT controls. All compounds were tested at a concentration of 10 µM, a dose commonly used in zebrafish studies as a compromise between efficacy and toxicity for small molecules, consistent with our previous dose–response observations.

Within the MAPK Inhibitor Library, 149 compounds were evaluated. Of these, 23 compounds significantly increased the total distance traveled by *cdkl5^-/-^* larvae compared to DMSO (vehicle)-treated mutants. Notably, nine compounds (resveratrol, VX-702, GDC-0879, WHI-P154, quercitrin, VX-11e, shanzhiside methyl ester, pamapimod and LY3214996) restored locomotor activity to levels that were not statistically different from those of DMSO-treated WT larvae. A summary of the MAPK inhibitor screening results is shown in Figure 3.

**Figure 3.**
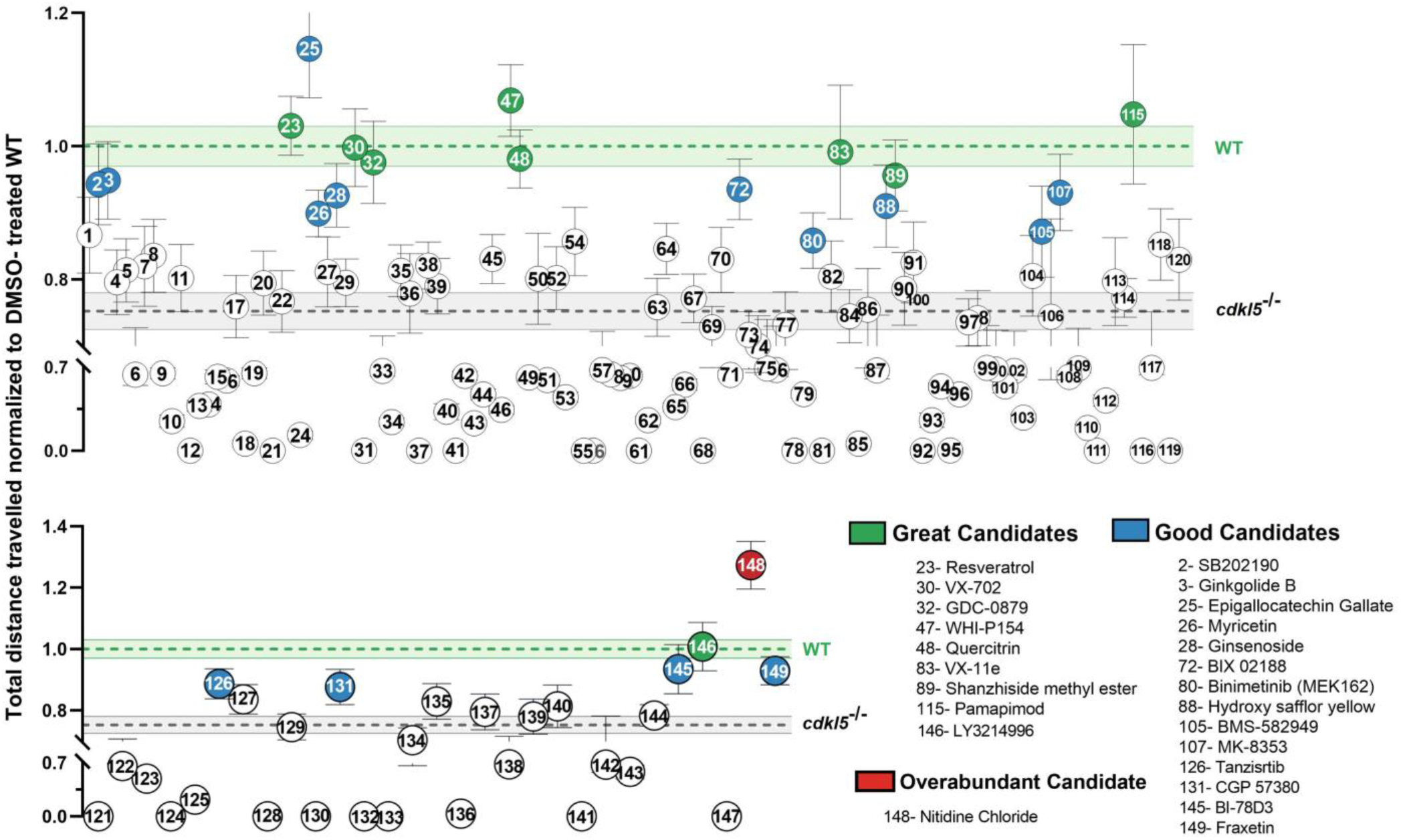
Screening of a MAPK Inhibitor Library for rescue of abnormal swimming behavior in *cdkl5*^⁻/⁻^ larvae. Total distance traveled by 5 dpf *cdkl5^-/-^* zebrafish larvae following treatment with 149 compounds from the MAPK Inhibitor Library (10 μM). Each dot represents the mean ± SEM distance traveled by 48 *cdkl5*^-/-^ larvae per compound, normalized to DMSO-treated WT controls. The green and grey horizontal lines indicate the mean ± SEM distance traveled by DMSO-treated WT and DMSO-treated *cdkl5^-/-^*larvae, respectively. Statistical analysis was performed using one-way ANOVA followed by Dunnett’s post hoc test. Compounds classified as great candidates (green dots) significantly increased (*p*<0.05) the distance traveled by *cdkl5*^⁻/⁻^ larvae to levels not significantly different from those of DMSO-treated WT controls (*p*>0.05). Compounds classified as good candidates (blue dots) significantly increased (*p*<0.05) the distance traveled by *cdkl5*^⁻/⁻^ larvae but remained significantly lower than that of DMSO-treated WT controls (*p*<0.05). The compound classified as overabundant candidate (red dot) resulted in a significantly greater distance traveled by *cdkl5*^⁻/⁻^ larvae compared to both DMSO-treated *cdkl5*^⁻/⁻^ and WT control larvae (*p*<0.05).

Within the Histone Modification Library, 21 compounds were tested. Seven of these compounds significantly increased the total distance traveled by *cdkl5^-/-^* larvae compared with DMSO-treated mutant larvae. Among these, three compounds (resveratrol, fisetin and AMI-1), restored locomotor activity to levels that were not statistically different from those of DMSO-treated WT larvae. A summary of the Histone Modification Library screening results is presented in Figure 4.

**Figure 4.**
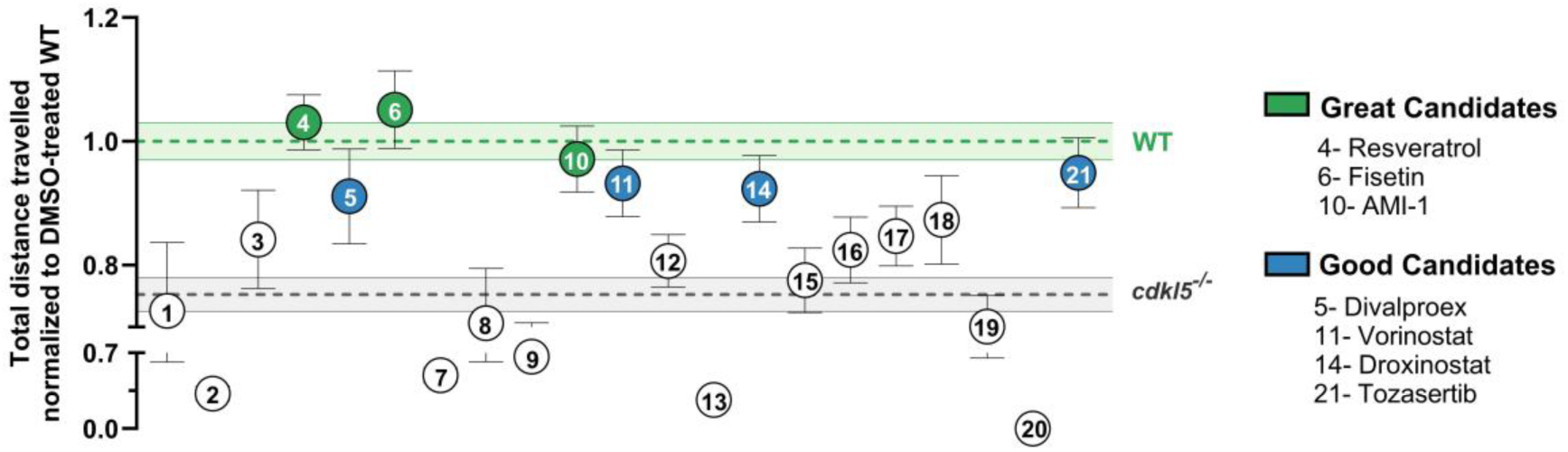
Screening of a Histone Modification Library for rescue of abnormal swimming behavior in *cdkl5*^⁻/⁻^ larvae. Total distance traveled by 5 dpf *cdkl5^-/-^* zebrafish larvae following treatment with 21 compounds from the Histone Modification Library (10 μM). Each dot represents the mean ± SEM distance traveled by 48 *cdkl5*^-/-^ larvae per compound, normalized to DMSO-treated WT controls. The green and grey horizontal lines indicate the mean ± SEM distance traveled by DMSO-treated WT and DMSO-treated *cdkl5^-/-^* larvae, respectively. Statistical analysis was performed using one-way ANOVA followed by Dunnett’s post hoc test. Compounds classified as great candidates (green dots) significantly increased (*p*<0.05) the distance traveled by *cdkl5*^⁻/⁻^ larvae to levels not significantly different from those of DMSO-treated WT controls (*p*>0.05). Compounds classified as good candidates (blue dots) significantly increased (*p*<0.05) the distance traveled by *cdkl5*^⁻/⁻^ larvae but remained significantly lower than that of DMSO-treated WT controls (*p*<0.05).

To investigate whether the observed rescue on swimming behavior was *cdkl5*-specific rather than due to a general locomotor stimulant effect, WT larvae were treated with 10 µM of the candidate compounds, and total distance traveled was quantified (Figure 5). A subset of four compounds was selected for this and subsequent in-depth analyses: fisetin, resveratrol and VX-702 from the group of great candidates, along with divalproex from the group of good candidates. Our results showed that the treatment of WT larvae with these compounds did not result in a significant difference in total distance traveled compared to DMSO-treated WT larvae (Figure 5). Overall, these findings indicate that the selected candidate compounds do not affect baseline locomotor activity in WT zebrafish, indicating that the rescue observed in mutant fish is *cdkl5*-specific.

**Figure 5.**
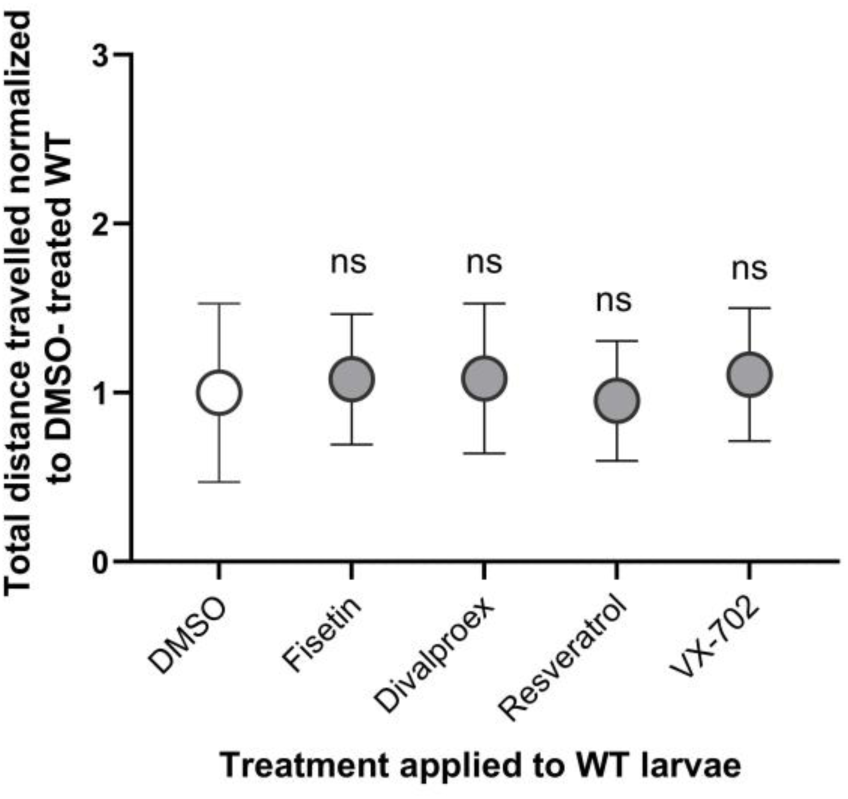
Effect of candidate molecules on the swimming behavior of WT larvae. The total distance traveled of 5 dpf WT larvae treated with 10 μM of candidate molecules was analyzed and compared with WT larvae treated with 0.1% DMSO (vehicle control). Each dot represents the mean ± SEM distance traveled by 48 WT larvae per compound, normalized to DMSO-treated WT controls. Statistical analysis was performed using the Kruskal–Wallis test followed by Dunn’s multiple comparisons test. No significant differences were detected between groups.

### 3.3. Effect of candidate compounds on craniofacial structures in *cdkl5^-/-^* mutant zebrafish

Previous studies have reported that *cdkl5*^-/-^ zebrafish also display craniofacial defects, indicating that Cdkl5 deficiency affects not only neuronal function but also cartilage developmental processes^27–29,34^. To determine whether candidate compounds identified through behavioral screening could also ameliorate these morphological defects, we evaluated their effects on craniofacial cartilage formation in *cdkl5^-/-^*larvae. For this end, *cdkl5^-/-^* mutants were treated from 3 to 5 dpf with fisetin, divalproex, resveratrol and VX-702, after which craniofacial cartilage structures were visualized by alcian blue staining (Figure 6A). Our results confirmed the altered craniofacial morphology of *cdkl5*^-/-^ larvae. Quantitative morphometric analysis (Figure 6B) revealed significant defects in multiple cartilage parameters, including a reduction in palatoquadrate length (Pq-l) and decreased ceratohyal length (Ch-l) and width (Ch-w) (Figure 6C). All tested compounds significantly increased Pq-l in *cdkl5*^-/-^ larvae, although not to WT levels. Similarly, fisetin, divalproex and resveratrol produced a partial rescue of Ch-l, whereas VX-702 did not exert a significant effect on this parameter. In contrast, a significant increase in Ch-w was observed only in *cdkl5*^-/-^ larvae treated with fisetin or VX-702, however, values remained significantly lower than those of WT larvae (Figure 6C). Overall, none of the tested compounds fully rescued all craniofacial parameters, indicating partial and parameter-specific effects.

**Figure 6.**
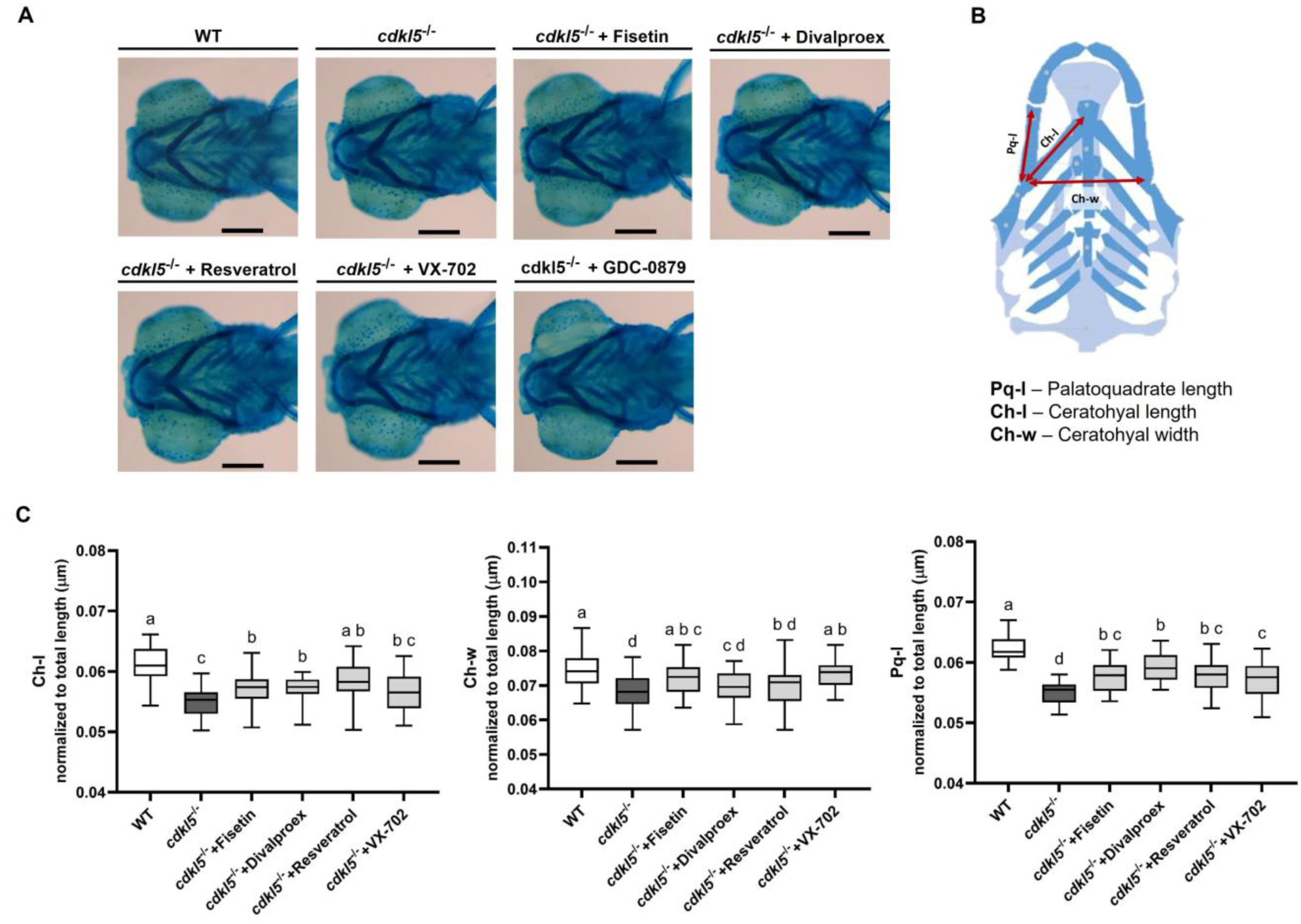
Effect of candidate molecules on craniofacial cartilage structures in 5 dpf *cdkl5^-/-^* mutant zebrafish larvae. **A)** Representative images of the WT, DMSO-treated *cdkl5*^⁻/⁻^ larvae and *cdkl5^-/-^* larvae following drug treatment with fisetin, divalproex, resveratrol and VX-702, stained with Alcian blue. **B)** Schematic illustration of the analyzed craniofacial cartilage parameters, adapted from Raterman *et al.*^89^ **C)** Quantification of craniofacial cartilage parameters, including ceratohyal cartilage length (Ch-l), palatoquadrate length (Pq-l), and ceratohyal cartilage width (Ch-w). Data are presented as median with interquartile range (n=29). Statistical analysis was performed using one-way ANOVA followed by Tukey’s multiple comparisons test. Significant differences are indicated by distinct lowercase letters.

### 3.4. Capability of candidate compounds to rescue gene expression defects in *cdkl5^-/-^* mutant zebrafish

In addition to behavioral and morphological abnormalities, Cdkl5 deficiency has been associated with dysregulation affecting genes involved in neuronal function, cytoskeletal organization, and extracellular matrix composition^29^. To determine whether selected compounds identified through behavioral screening could also restore molecular defects associated with *cdkl5* loss, we examined the effects of these molecules on the expression of genes previously reported to be downregulated in *cdkl5*⁻^/^⁻ larvae. Quantitative real-time PCR analysis was performed on *cdkl5*⁻^/^⁻ larvae treated with fisetin, divalproex, resveratrol and VX-702, focusing on the expression of *arhgef2*, *iqgap1*, *ahsa1b*, and *col1a1b* and compared with those of untreated *cdkl5*⁻^/^⁻ mutant larvae and WT siblings (Figure 7).

**Figure 7.**
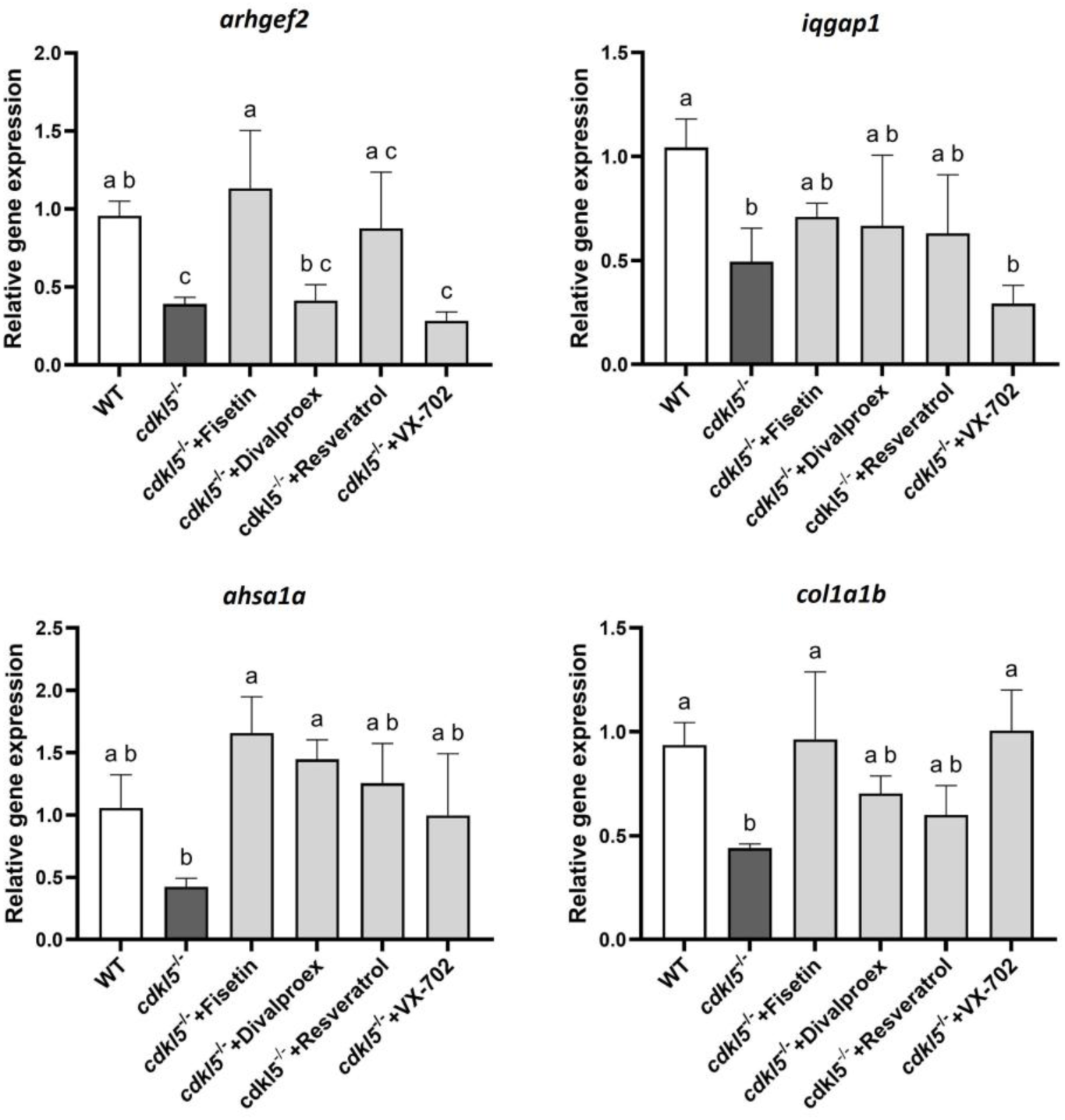
Gene expression analysis in 5 dpf *cdkl5*^-/-^ larvae following treatment with candidate molecules. RT-qPCR analysis was performed on larvae treated with 10 µM of fisetin, divalproex, resveratrol or VX-702 for 48h. WT and *cdkl5*^-/-^ control groups were treated with 0.1% DMSO as vehicle. Relative gene expression levels are presented as mean±SD of three biological replicates, except for the resveratrol group, which included two biological replicates. Each replicate consisted of a pool of 15 larvae. Statistical analysis was performed using one-way ANOVA followed by Tukey’s multiple comparisons test. Statistically significant differences are indicated by distinct lowercase letters.

Treatment with fisetin resulted in a significant upregulation of *arhgef2*, *ahsa1b*, and *col1a1b* in *cdkl5*^⁻/⁻^ larvae, restoring their expression to levels that were not statistically different from those observed in WT controls (Figure 7). Treatment with divalproex also resulted in a significant increase in *ahsa1b* expression, restoring levels comparable to those observed in WT larvae. However, no significant changes were detected in the expression of *arhgef2*, *iqgap1*, or *col1a1b* following divalproex treatment (Figure 7). Treatment with VX-702 significantly increased *col1a1b* expression in *cdkl5*^-/-^ mutants, restoring levels comparable to those observed in WT. In contrast, resveratrol treatment did not produce significant changes in the expression of any of the genes analyzed in *cdkl5*^-/-^ larvae (Figure 7). Overall, these results demonstrate that the selected candidate compounds can differentially affect/rescue gene expression alterations associated with *cdkl5* deficiency *in vivo*.

## 4. DISCUSSION

Zebrafish larvae have become one of the most widely used *in vivo* models for high-throughput drug screening in recent years^35,36^. It offers experimental advantages that enable rapid assessment of therapeutic compounds in a broad spectrum of disease models, including those affecting neurodevelopment, which is difficult to achieve using conventional rodent models^26,35,37^. Applying zebrafish-based screening to CDKL5 deficiency disorder (CDD), a severe condition for which effective treatments remain scarce^15^, provides a powerful strategy for identifying candidate drugs capable of rescuing functional deficits. Here, we report a locomotor behavior–based drug screening assay in a zebrafish model of CDD that, to the best of our knowledge, has not previously been performed in a preclinical animal model of this disorder. Consistent with key features of the human condition, *cdkl5*^⁻/⁻^ larvae exhibit motor impairments, including reduced swimming activity and shortened motor neurons^27–29^, which provide a disease-relevant functional readout for therapeutic assessment. Locomotor behavior, quantified as distance traveled, was selected as the primary endpoint because it offers a rapid, simple, and high-throughput measure while providing a robust evaluation of motor function^38,39^. This screen identified small molecules that improved impaired locomotor activity in 5 dpf *cdkl5*^⁻/⁻^ larvae following two days of treatment. Drug treatment was initiated at 3 dpf, a developmental stage at which embryos have naturally exited the chorion, thereby avoiding the need for manual dechorionation or enzymatic treatments that could interfere with subsequent analyses. Behavioral assessment was performed at 5 dpf, prior to the onset of feeding, to minimize variability that could influence locomotor behavior and the evaluation of drug effects. At this stage, larvae display robust and reproducible swimming behavior, enabling reliable redout of locomotor deficits and drug-induced rescue.

Prior to screening our libraries, we evaluated four previously identified compounds using our locomotor assay in the *cdkl5* zebrafish model: ivabradine, a blocker of hyperpolarization-activated cyclic nucleotide-gated (HCN) channels; solifenacin, an antagonist of muscarinic acetylcholine receptors; AZD1080, an inhibitor of glycogen synthase kinase-3 (GSK3); and crenigacestat, a γ-secretase inhibitor that plays a crucial role in Notch signaling. Negraes et al. reported that these molecules were able to rescue the hyperexcitability and outward radial cellular migration defects in patient-derived CDD spheroids^33^. Consistent with these findings, our *in vivo* zebrafish model revealed a significant improvement in locomotor behavior following treatment with solifenacin, ivabradine and crenigacestat. The complete rescue achieved with solifenacin suggests a role for cholinergic signaling in CDKL5 deficiency and its modulation may represent a promising therapeutic strategy. Recent evidence shows that CDKL5 directly phosphorylates the voltage-gated calcium channel Cav2.3, a downstream target of muscarinic acetylcholine receptors^40^. Loss of this phosphorylation in *Cdkl5*-deficient mice results in enhanced channel activity due to delayed inactivation and increased cholinergic stimulation, thereby elevating neuronal excitability and contributing to behavioral deficits and increased susceptibility to epilepsy^40,41^. Pharmacological blockage of muscarinic receptors with solifenacin may therefore mitigate this pathological gain-of-function, reduce aberrant network activity, and promote normalization of motor circuit function. In contrast, AZD1080 did not ameliorate the locomotor impairment observed in *cdkl5*^⁻/⁻^ larvae, at least under the present experimental conditions. One possible explanation is that the previously reported benefits were identified in a cell-based system, which do not fully capture the complexity of neural circuits present in whole organisms such as zebrafish. Alternatively, the behavioral readout assessed here may depend on mechanisms that are distinct from, or less sensitive to, those modulated by GSK3 inhibition.

In 2022, ganaxolone, a positive allosteric modulator of GABA_A_ receptors, became the first and only approved drug for treatment of CDD-associated seizures, following a phase 3 trial demonstrating a ∼30.7% reduction in seizure frequency in affected individuals^16^. Based on this clinical efficacy, we investigated whether ganaxolone could also ameliorate the locomotor behavior of *cdkl5*^⁻/⁻^ larvae. No significant improvement was observed, suggesting that seizure suppression and restoration of motor behavior may depend on distinct circuit mechanisms. The locomotor deficits captured in our assay may therefore result from abnormalities that are not corrected by potentiation of GABAergic signaling alone. Given that this zebrafish mutant presents higher susceptibility to seizures than WT siblings following PTZ induction^27,28^, it would be important to determine whether ganaxolone is also capable of reducing seizure activity in this zebrafish model.

The MAPK pathway regulates essential cellular processes, including proliferation, differentiation, and apoptosis, and its dysregulation contributes to the pathogenesis of numerous conditions, including neurodevelopmental and neurodegenerative disorders^42–44^. Notably, dysregulation of MAPK/ERK pathway has been reported in CDKL5 mouse and cell models, where loss of CDKL5 results in impaired phosphorylation levels of extracellular signal-regulated kinase (ERK1/2)^45–48^. Furthermore, CDKL5 has been implicated in epigenetic regulation through its ability to phosphorylate multiple proteins involved in chromatin remodeling and DNA methylation, including histone deacetylase 4 (HDAC4)^49^, E1A-Binding Protein P400 (EP400)^50^, GATA zinc finger domain containing 2A (GATAD2A)^51^, methyl-CpG binding protein 2 (MeCP2)^52^, and DNA methyltransferase 1 (DNMT1)^53^. Accumulating evidence indicates that HDACs play crucial roles in brain development and neuron survival. Their dysfunction has been associated with the pathogenesis of a broad range of disorders, including neurological diseases. On this basis, pharmacological inhibitors of HDAC activity are increasingly being investigated as a potential therapeutic strategy for these conditions^54–56^. Interestingly, loss of CDKL5 function in a knockout mouse model resulted in reduced HDAC4 phosphorylation, promoting its nuclear translocation and concomitant decrease in histone acetylation, ultimately leading to altered gene expression. These changes triggered apoptosis of newborn granule neurons and impaired dendritic development and synaptic organization of hippocampal neurons. Notably, these defects were rescued upon inhibition of HDAC4 activity^49^. Given the growing evidence demonstrating a direct interaction and regulatory role of CDKL5 in the MAPK signaling pathway and chromatin remodeling, we chose the MAPK Inhibitor and Histone Modification Libraries for our screening assay.

From the initial set of 170 molecules screened using our platform, we identified 18 and 12 small molecules that partially or fully rescued the locomotor deficits of *cdkl5*^⁻/⁻^ mutant larvae, respectively, as determined by an increase in total distance traveled. Among those, ten were natural biological compounds, including antioxidants and anti-inflammatory agents, with a broad spectrum of activity (resveratrol, fisetin, quercitrin, shanzhiside methyl ester, myricitrin, ginkgolide b, ginsenosides, hydroxysafflor yellow A, fraxetin and epigallocatechin gallate). Additionally, six (GDC-0879, Binimetinib, VX-11e, MK-8353, LY3214996 and BIX 02188) and five (SB202190, VX-702, BMS- 582949, Pamapimod and Tanzisertib) direct MAPK antagonists that specifically target the RAF–MEK–ERK axis and p38 cascade, respectively, also demonstrated partial or complete locomotor rescue. A similar result was obtained with three HDAC inhibitors (vorinostat, droxinostat and divalproex sodium) and one histone methyltransferase inhibitor (AMI-1).

A subset of four molecules (fisetin, divalproex, resveratrol, and VX-702) was selected for in-depth analysis. These compounds were prioritized based on (i) their ability to rescue locomotor behavior in *cdkl5*^⁻/⁻^ mutant larvae, (ii) published evidence supporting their activity in biological systems relevant to CDKL5 deficiency disorder (neuronal, neuromuscular, and skeletal systems; Table 2), and (iii) their commercial availability from suppliers independent of the original screening library. Although divalproex did not fully rescue the locomotor phenotype of *cdkl5*^⁻/⁻^ mutants, it was included in subsequent analyses because it is an FDA-approved anticonvulsant widely used for seizure management^57,58^. It is composed of sodium valproate and valproic acid (1:1 molar ratio) that have been reported to improve motor function in certain animal models of neuromuscular dysfunction^59–61^. Fisetin and resveratrol were of particular interest due to multiple reports describing their anticonvulsant and neuroprotective effects in zebrafish and rodent models of seizures^62–67^. In addition, both compounds have been associated with enhanced motor neuron survival and function^68–74^ and modulation of skeletal developmental or craniofacial abnormalities. In addition, both compounds have been associated with enhanced motor neuron survival and function^68–74^ and modulation of skeletal developmental or craniofacial abnormalities^75–81^. Importantly, treatment of WT larvae with fisetin, divalproex, resveratrol, or VX-702 did not alter baseline locomotor activity, indicating that the behavioral rescue observed in *cdkl5*^⁻/⁻^ mutants is unlikely to result from nonspecific locomotor stimulation. This genotype-selective effect strengthens the conclusion that these compounds are likely acting on pathways disrupted by *cdkl5* loss, correcting molecular or cellular dysfunctions associated with CDKL5 deficiency rather than by broadly enhancing motor output. Such genotype-dependent rescue is particularly relevant for translational development, as it indicates therapeutic potential without perturbing physiological motor function.

**Table 2.**
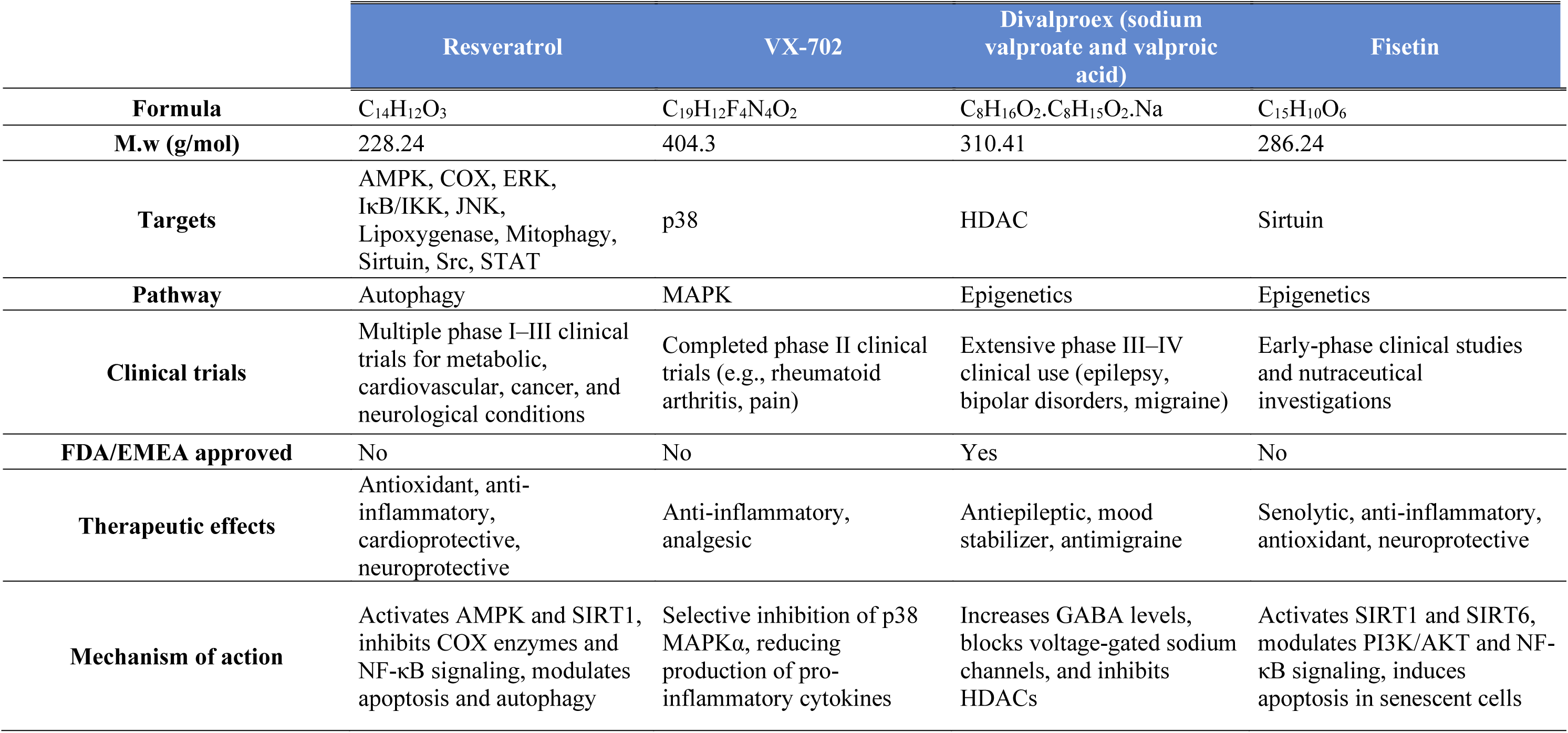
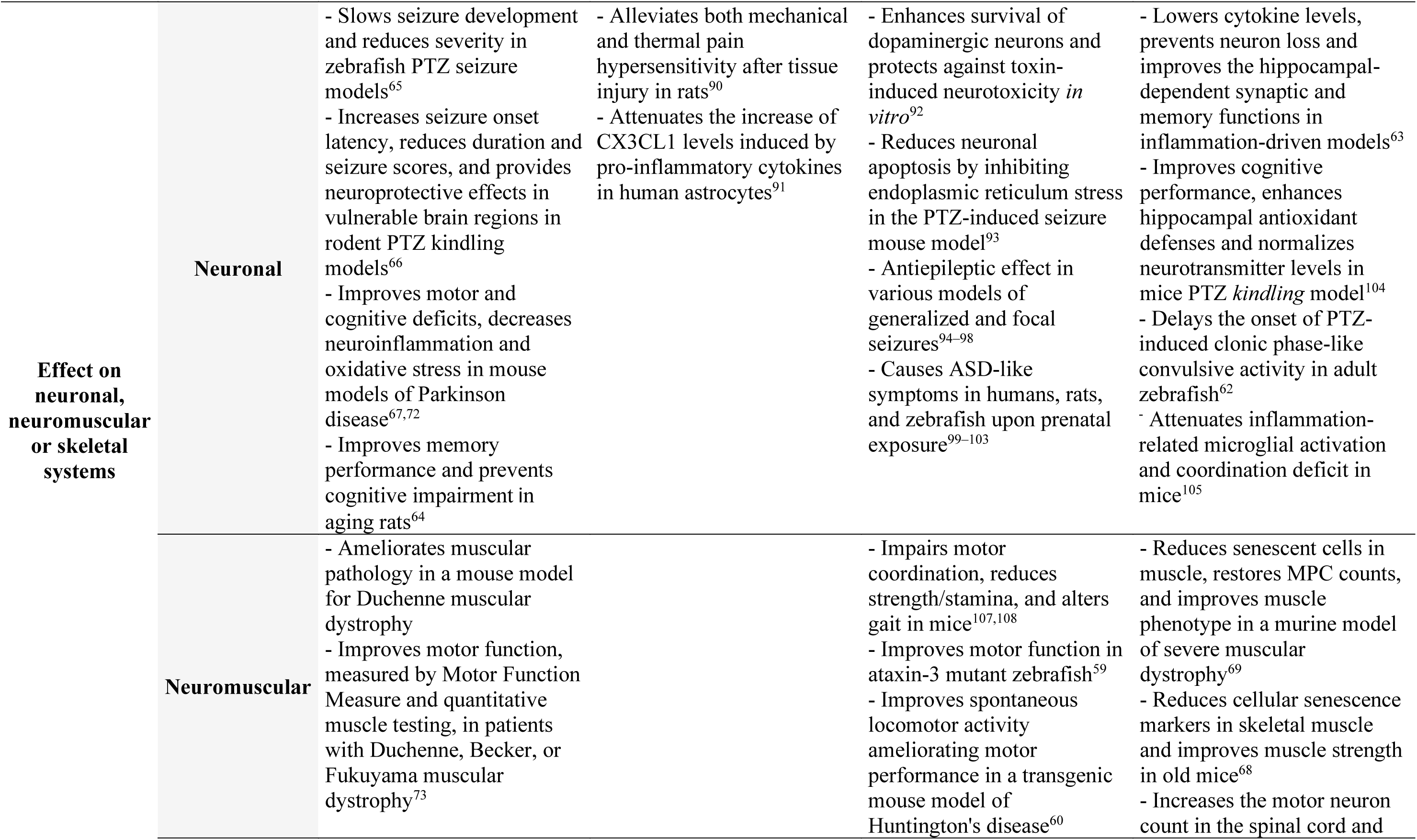
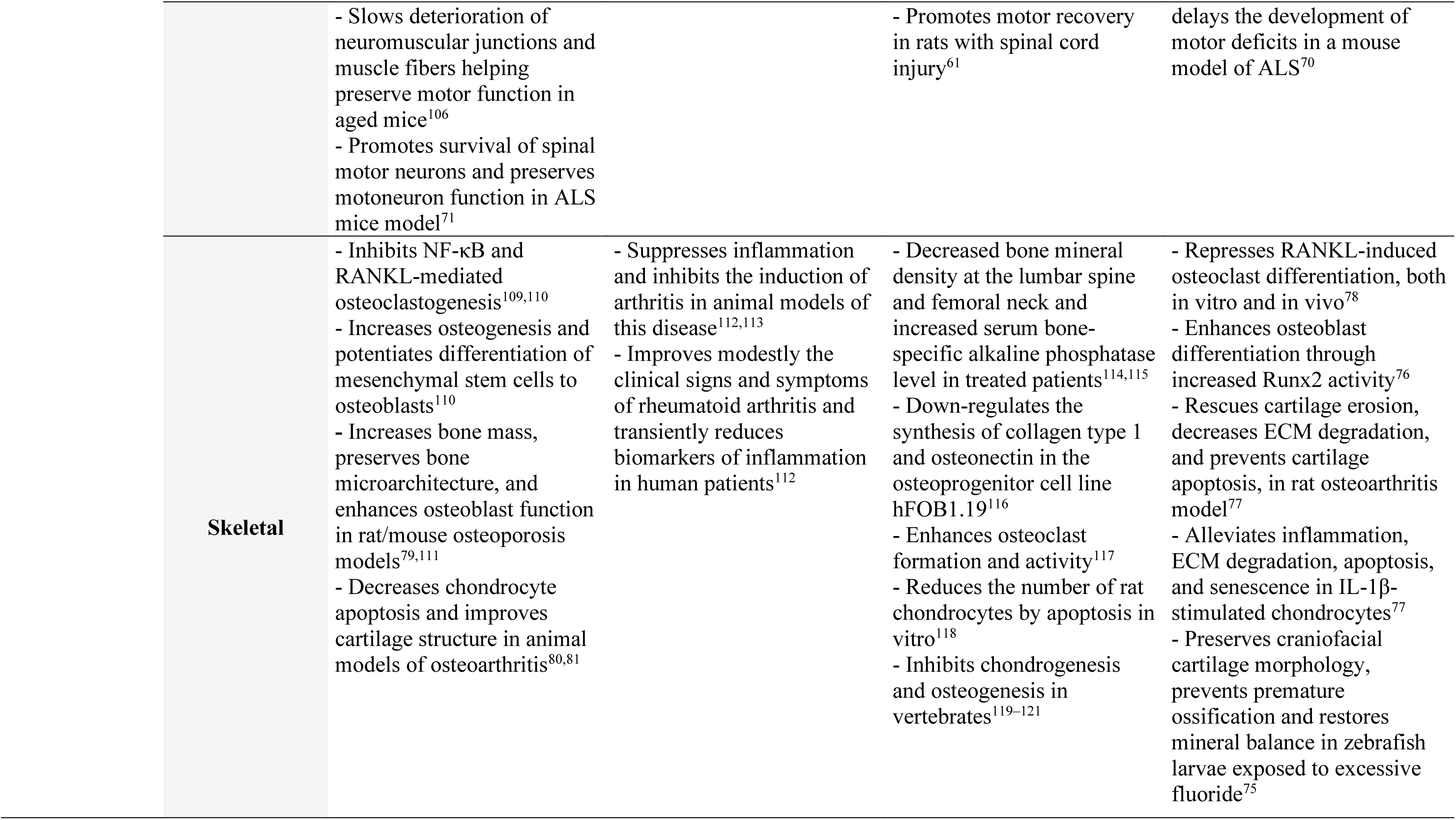
Summary of chemical properties, mechanisms of action, and reported effects of selected compounds on neuronal, neuromuscular or skeletal systems.

|  | Resveratrol | VX-702 | Divalproex (sodium valproate and valproic acid) | Fisetin |
| --- | --- | --- | --- | --- |
| **Formula** | C<sub>14</sub>H<sub>12</sub>O<sub>3</sub> | C<sub>19</sub>H<sub>12</sub>F<sub>4</sub>N<sub>4</sub>O<sub>2</sub> | C<sub>8</sub>H<sub>16</sub>O<sub>2</sub>.C<sub>8</sub>H<sub>15</sub>O<sub>2</sub>.Na | C<sub>15</sub>H<sub>10</sub>O<sub>6</sub> |
| **M.w (g/mol)** | 228.24 | 404.3 | 310.41 | 286.24 |
| **Targets** | AMPK, COX, ERK, IκB/IKK, JNK, Lipoxygenase, Mitophagy, Sirtuin, Src, STAT | p38 | HDAC | Sirtuin |
| **Pathway** | Autophagy | MAPK | Epigenetics | Epigenetics |
| **Clinical trials** | Multiple phase I–III clinical trials for metabolic, cardiovascular, cancer, and neurological conditions | Completed phase II clinical trials (e.g., rheumatoid arthritis, pain) | Extensive phase III–IV clinical use (epilepsy, bipolar disorders, migraine) | Early-phase clinical studies and nutraceutical investigations |
| **FDA/EMA approved** | No | No | Yes | No |
| **Therapeutic effects** | Antioxidant, anti-inflammatory, cardioprotective, neuroprotective | Anti-inflammatory, analgesic | Antiepileptic, mood stabilizer, antimigraine | Senolytic, anti-inflammatory, antioxidant, neuroprotective |
| **Mechanism of action** | Activates AMPK and SIRT1, inhibits COX enzymes and NF-κB signaling, modulates apoptosis and autophagy | Selective inhibition of p38 MAPKα, reducing production of pro-inflammatory cytokines | Increases GABA levels, blocks voltage-gated sodium channels, and inhibits HDACs | Activates SIRT1 and SIRT6, modulates PI3K/AKT and NF-κB signaling, induces apoptosis in senescent cells |

|  |  |  |  |  |  |
| --- | --- | --- | --- | --- | --- |
| **Effect on neuronal, neuromuscular or skeletal systems** | **Neuronal** | - Slows seizure development and reduces severity in zebrafish PTZ seizure models<sup>65</sup><br>- Increases seizure onset latency, reduces duration and seizure scores, and provides neuroprotective effects in vulnerable brain regions in rodent PTZ kindling models<sup>66</sup><br>- Improves motor and cognitive deficits, decreases neuroinflammation and oxidative stress in mouse models of Parkinson disease<sup>67,72</sup><br>- Improves memory performance and prevents cognitive impairment in aging rats<sup>64</sup> | - Alleviates both mechanical and thermal pain hypersensitivity after tissue injury in rats<sup>90</sup><br>- Attenuates the increase of CX3CL1 levels induced by pro-inflammatory cytokines in human astrocytes<sup>91</sup> | - Enhances survival of dopaminergic neurons and protects against toxin-induced neurotoxicity *in vitro*<sup>92</sup><br>- Reduces neuronal apoptosis by inhibiting endoplasmic reticulum stress in the PTZ-induced seizure mouse model<sup>93</sup><br>- Antiepileptic effect in various models of generalized and focal seizures<sup>94–98</sup><br>- Causes ASD-like symptoms in humans, rats, and zebrafish upon prenatal exposure<sup>99–103</sup> | - Lowers cytokine levels, prevents neuron loss and improves the hippocampal-dependent synaptic and memory functions in inflammation-driven models<sup>63</sup><br>- Improves cognitive performance, enhances hippocampal antioxidant defenses and normalizes neurotransmitter levels in mice PTZ *kindling* model<sup>104</sup><br>- Delays the onset of PTZ-induced clonic phase-like convulsive activity in adult zebrafish<sup>62</sup><br>- Attenuates inflammation-related microglial activation and coordination deficit in mice<sup>105</sup> |
|  | **Neuromuscular** | - Ameliorates muscular pathology in a mouse model for Duchenne muscular dystrophy<br>- Improves motor function, measured by Motor Function Measure and quantitative muscle testing, in patients with Duchenne, Becker, or Fukuyama muscular dystrophy<sup>73</sup> |  | - Impairs motor coordination, reduces strength/stamina, and alters gait in mice<sup>107,108</sup><br>- Improves motor function in ataxin-3 mutant zebrafish<sup>59</sup><br>- Improves spontaneous locomotor activity ameliorating motor performance in a transgenic mouse model of Huntington's disease<sup>60</sup> | - Reduces senescent cells in muscle, restores MPC counts, and improves muscle phenotype in a murine model of severe muscular dystrophy<sup>69</sup><br>- Reduces cellular senescence markers in skeletal muscle and improves muscle strength in old mice<sup>68</sup><br>- Increases the motor neuron count in the spinal cord and |
|  | - Slows deterioration of neuromuscular junctions and muscle fibers helping preserve motor function in aged mice<sup>106</sup><br>- Promotes survival of spinal motor neurons and preserves motoneuron function in ALS mice model<sup>71</sup> |  | - Promotes motor recovery in rats with spinal cord injury<sup>61</sup> | delays the development of motor deficits in a mouse model of ALS<sup>70</sup> |
| **Skeletal** | - Inhibits NF-κB and RANKL-mediated osteoclastogenesis<sup>109,110</sup><br>- Increases osteogenesis and potentiates differentiation of mesenchymal stem cells to osteoblasts<sup>110</sup><br>- Increases bone mass, preserves bone microarchitecture, and enhances osteoblast function in rat/mouse osteoporosis models<sup>79,111</sup><br>- Decreases chondrocyte apoptosis and improves cartilage structure in animal models of osteoarthritis<sup>80,81</sup> | - Suppresses inflammation and inhibits the induction of arthritis in animal models of this disease<sup>112,113</sup><br>- Improves modestly the clinical signs and symptoms of rheumatoid arthritis and transiently reduces biomarkers of inflammation in human patients<sup>112</sup> | - Decreased bone mineral density at the lumbar spine and femoral neck and increased serum bone-specific alkaline phosphatase level in treated patients<sup>114,115</sup><br>- Down-regulates the synthesis of collagen type 1 and osteonectin in the osteoprogenitor cell line hFOB1.19<sup>116</sup><br>- Enhances osteoclast formation and activity<sup>117</sup><br>- Reduces the number of rat chondrocytes by apoptosis in vitro<sup>118</sup><br>- Inhibits chondrogenesis and osteogenesis in vertebrates<sup>119-121</sup> | - Represses RANKL-induced osteoclast differentiation, both in vitro and in vivo<sup>78</sup><br>- Enhances osteoblast differentiation through increased Runx2 activity<sup>76</sup><br>- Rescues cartilage erosion, decreases ECM degradation, and prevents cartilage apoptosis, in rat osteoarthritis model<sup>77</sup><br>- Alleviates inflammation, ECM degradation, apoptosis, and senescence in IL-1β-stimulated chondrocytes<sup>77</sup><br>- Preserves craniofacial cartilage morphology, prevents premature ossification and restores mineral balance in zebrafish larvae exposed to excessive fluoride<sup>75</sup> |

Cdkl5 deficiency impacts not only neuronal and motor function but also cartilage development and craniofacial morphogenesis^27–29,34^. Although none of the tested compounds fully restored craniofacial architecture, all produced partial recovery of specific structural parameters, with fisetin showing the most consistent and pronounced effects across multiple measurements. These findings suggest that the molecular pathways underlying locomotor impairment may partially overlap with those regulating craniofacial development. However, the incomplete rescue observed across craniofacial parameters highlights the complexity of Cdkl5-dependent processes and indicates that multiple dysregulated pathways may contribute to the morphological phenotype. Cdkl5 deficiency has been linked to widespread transcriptional alterations impacting genes that regulate cytoskeletal dynamics, neuronal signaling, and extracellular matrix organization^29,82^. In this study, we examined whether the selected compounds could also restore molecular alterations previously reported in *cdkl5*^⁻/⁻^ larvae. Specifically, we analyzed *arhgef2*, which encodes a Rho guanine nucleotide exchange factor, a direct CDKL5 target implicated in microtubule stability, dendritic spine maintenance, and neuronal polarity^83^; *iqgap1*, encoding a scaffold protein that regulates actin dynamics and forms a complex with CDKL5 to promote dendritic arborization^34^; *ahsa1a*, a co-chaperone of Hsp90 implicated in protein folding and cellular stress responses^84,85^; and *col1a1b*, encoding the α1 chain of type I collagen, a key structural component of the extracellular matrix required for skeletal morphogenesis^86–88^. Fisetin treatment significantly upregulated *arhgef2*, *ahsa1a*, and *col1a1b* expression in *cdkl5*^⁻/⁻^ larvae, restoring transcript levels to those comparable with WT controls, suggesting broad molecular modulation across cytoskeletal, stress-response, and extracellular matrix pathways. Divalproex restored *ahsa1a* expression but did not affect *arhgef2*, *iqgap1*, or *col1a1b*, indicating a more selective transcriptional effect, potentially related to its known epigenetic and histone deacetylase–inhibitory activity. VX-702 significantly increased *col1a1b* expression, consistent with its partial rescue of craniofacial parameters and suggesting a link between p38 MAPK modulation and extracellular matrix regulation in Cdkl5 deficiency. In contrast, resveratrol did not significantly alter the expression of the genes analyzed, despite improving behavioral and craniofacial parameters. This finding may suggest that resveratrol does not directly modulate the pathways regulating these genes or that it exerts its effects through mechanisms not reflected at the transcriptional level of these targets, such as post-transcriptional regulation or functional modulation. However, it is important to note that gene expression analysis for the resveratrol-treated group was performed with only two biological replicates. Therefore, these results should be interpreted cautiously and require further validation with increased sample size.

While this study demonstrates the utility of the zebrafish *cdkl5* mutant as a platform for phenotypic drug screening, some limitations should be acknowledged. First, the use of a single concentration (10 µM) administered over a relatively short treatment period of 48 hours may have resulted in the exclusion of potentially effective compounds that did not exhibit detectable activity under these specific conditions but might demonstrate efficacy at different concentrations or with prolonged exposure. This approach reflected the need to ensure screening throughput and cost-effectiveness while selecting a concentration commonly used and reported as active in zebrafish *in vivo* screening studies. Second, zebrafish larvae represent an early developmental stage, and drug effects observed at this stage may not directly translate to later developmental stages or to mammalian systems. Finally, motor impairment represents only one component of the complex CDD phenotype. Given that drug-resistant seizures are a major clinical feature of CDD, the integration of complementary assays, such as calcium imaging and electrophysiology recordings, would strengthen the evaluation of therapeutic candidates. Future studies should focus on elucidating the molecular mechanisms underlying the effects of these molecules by comprehensively characterizing their targets within the signaling networks disrupted by *cdkl5* deficiency. High-throughput approaches such as transcriptomic profiling (RNA-seq) and proteomic analyses will be particularly valuable for identifying differentially regulated genes, signaling pathways, and protein interaction networks associated with phenotypic rescue.

In conclusion, our study establishes a simple and rapid zebrafish-based phenotypic screening platform and highlights its utility for the discovery of novel compounds for CDD. This approach enabled the identification of small molecules capable of partially or fully rescuing key phenotypic alterations in the *cdkl5* mutant and may provide a promising direction for the development of new therapeutic strategies for CDD. Validation in complementary models, including rodent systems and patient-derived cellular models, will be essential to determine the translational potential of the identified candidates.

## Acknowledgements

This work was supported by the Million Dollar Bike Grant program from the Orphan Disease Center, University of Pennsylvania, USA (Award Number: MDBR-19-104-CDKL5) and a research grant from the University of Pennsylvania Orphan Disease Center (CDKL5-22-103-02) on behalf of the Loulou Foundation. And received Portuguese national funds from FCT - Foundation for Science and Technology through the project Zfscreen (2022.06526.PTDC) and contracts UID/04326/2025, UID/PRR/04326/2025 and LA/P/0101/2020 (DOI:10.54499/LA/P/0101/2020), and from the operational programs CRESC Algarve 2020 and COMPETE 2020 through contract EMBRC.PT ALG-01-0145-FEDER-022121. Tatiana Varela (https://doi.org/10.54499/SFRH/BD/144230/2019) and Débora Varela (SFRH/BD/141918/2018) were recipients of Ph.D. fellowships from FCT.

## Data availability statement

The data generated and analyzed during this study are available from the corresponding authors upon reasonable request.

## Author contributions

Conceptualization, MLC and NC; Formal analysis, TV, JS, DV, MD; Funding acquisition, MLC and NC; Investigation, TV, DV, JS, MD, AH, VP; Methodology, TV, DV, JS; Project administration, MLC; Resources, MLC and NC; Supervision, NC and MLC; Writing – original draft, TV; Writing – review & editing, TV, DV, NC and MLC.

## REFERENCES

1. The Lancet Global Health. The landscape for rare diseases in 2024. Lancet Glob. Health 12, e341 (2024).

2. Vavassori, S., Russell, S., Scotti, C. & Benvenuti, S. Unlocking the full potential of rare disease drug development: exploring the not-for-profit sector’s contributions to drug development and access. Front. Pharmacol. 15, 1441807 (2024).

3. Sun, W., Zheng, W. & Simeonov, A. Drug discovery and development for rare genetic disorders. Am. J. Med. Genet. A 173, 2307–2322 (2017).

4. Kaufmann, P., Pariser, A. R. & Austin, C. From scientific discovery to treatments for rare diseases - The view from the National Center for Advancing Translational Sciences - Office of Rare Diseases Research. Orphanet J. Rare Dis. 13, 196 (2018).

5. Wang, G. et al. Rare and undiagnosed diseases: From disease-causing gene identification to mechanism elucidation. Fundam. Res. 2, 918–928 (2022).

6. Jakimiec, M., Paprocka, J. & Śmigiel, R. CDKL5 deficiency disorder-A complex epileptic encephalopathy. Brain Sci. 10, 107 (2020).

7. Fehr, S. et al. The CDKL5 disorder is an independent clinical entity associated with early-onset encephalopathy. Eur J Hum Genet 21, 266–273 (2013).

8. Olson, H. E. et al. Cyclin-dependent kinase-Like 5 deficiency disorder: Clinical review. Pediatr. Neurol. 97, 18–25 (2019).

9. Bahi-Buisson, N. et al. Key clinical features to identify girls with CDKL5 mutations. Brain 131, 2647–2661 (2008).

10. Mangatt, M. et al. Prevalence and onset of comorbidities in the CDKL5 disorder differ from Rett syndrome. Orphanet J. Rare Dis. 11, 39 (2016).

11. Olson, H. E. et al. Current neurologic treatment and emerging therapies in CDKL5 deficiency disorder. J. Neurodev. Disord. 13, (2021).

12. Leonard, H. et al. CDKL5 deficiency disorder: clinical features, diagnosis, and management. Lancet Neurol. 21, 563–576 (2022).

13. Kalinowska-Doman, A., Strzelczyk, A. & Paprocka, J. Antiseizure medications in CDKL5 encephalopathy– systematic review. Seizure 131, 391–396 (2025).

14. Müller, A. et al. Retrospective evaluation of low long-term efficacy of antiepileptic drugs and ketogenic diet in 39 patients with CDKL5-related epilepsy. Eur. J. of Paediatr. Neurol. 20, 147–151 (2016).

15. Dell’Isola, G. B. et al. Current overview of CDKL-5 deficiency disorder treatment. Pediatr. Rep. 16, 21–25 (2024).

16. Pestana Knight, E. M., et al. Safety and efficacy of ganaxolone in patients with CDKL5 deficiency disorder: results from the double-blind phase of a randomised, placebo-controlled, phase 3 trial. Lancet Neurol. 21, 417–444 (2022).

17. Olson, H. E. et al. Long-term treatment with ganaxolone for seizures associated with cyclin-dependent kinase-like 5 deficiency disorder: Two-year open-label extension follow-up. Epilepsia 65, 37–45 (2024).

18. Carter, R. B. et al. Characterization of the anticonvulsant properties of ganaxolone (CCD 1042; 3α-hydroxy-3β-methyl-5α-pregnan-20-one), a selective, high-affinity, steroid modulator of the γ-aminobutyric acid (A) Receptor. J. Pharmacol. Exp. Ther. 280, 1284–1295 (1997).

19. Lattanzi, S., Riva, A. & Striano, P. Ganaxolone treatment for epilepsy patients: from pharmacology to place in therapy. Expert Rev. Neurother. 21, 1317–1332 (2021).

20. MacRae, C. A. & Peterson, R. T. Zebrafish as tools for drug discovery. Nat. Rev. Drug Discov. 14, 721–731 (2015).

21. Patton, E. E., Zon, L. I. & Langenau, D. M. Zebrafish disease models in drug discovery: from preclinical modelling to clinical trials. Nat. Rev. Drug Discov. 20, 611–628 (2021).

22. Zon, L. I. & Peterson, R. T. In vivo drug discovery in the zebrafish. Nat. Rev. Drug Discov. 4, 35–44 (2005).

23. Kari, G., Rodeck, U. & Dicker, A. P. Zebrafish: An emerging model system for human disease and drug discovery. Clin. Pharmacol. Ther. 82, 70–80 (2007).

24. MacRae, C. & Peterson, R. Zebrafish-based small molecule discovery. Chem. Biol. 10, 901–908 (2003).

25. Delvecchio, C., Tiefenbach, J. & Krause, H. M. The zebrafish: a powerful platform for in vivo, HTS drug discovery. Assay Drug Dev. Technol. 9, 354–361 (2011).

26. Dash, S. N. & Patnaik, L. Flight for fish in drug discovery: A review of zebrafish-based screening of molecules. Biol. Lett. 19, 20220541 (2023).

27. Varela, T., Varela, D., Martins, G., Conceição, N. & Cancela, M. L. Cdkl5 mutant zebrafish shows skeletal and neuronal alterations mimicking human CDKL5 deficiency disorder. Sci. Rep. 12, 9325 (2022).

28. Serrano, R. J. et al. Novel preclinical model for CDKL5 deficiency disorder. Dis. Model. Mech. 15, dmm049094 (2022).

29. Varela, T., Varela, D., Conceição, N. & Cancela, M. L. Transcriptomic profiling of zebrafish mutant for cdkl5 reveals dysregulated gene expression associated with neuronal, muscle, visual and skeletal development. Int. J. Mol. Sci. 26, 6069 (2025).

30. Busch-Nentwich, E. et al. Sanger Institute Zebrafish Mutation Project mutant data submission. ZFIN Direct Data Submission (2013).

31. Kimmel, C. B., Ballard, W. W., Kimmel, S. R., Ullmann, B. & Schilling, T. F. Stages of embryonic development of the zebrafish. Dev Dyn 203, 253–310 (1995).

32. Walker, M. B. & Kimmel, C. B. A two-color acid-free cartilage and bone stain for zebrafish larvae. Biotech. Histochem. 82, 23–28 (2007).

33. Negraes, P. D. et al. Altered network and rescue of human neurons derived from individuals with early-onset genetic epilepsy. Mol. Psychiatry 26, 7047–7068 (2021).

34. Barbiero, I., De Rosa, R. & Kilstrup-Nielsen, C. Microtubules: a key to understand and correct neuronal defects in CDKL5 deficiency disorder? Int. J. Mol. Sci. 20, 4075 (2019).

35. Lee, H. C., Lin, C. Y. & Tsai, H. J. Zebrafish, an in vivo platform to screen drugs and proteins for biomedical use. Pharmaceuticals 14, 500 (2021).

36. Rennekamp, A. J. & Peterson, R. T. 15 years of zebrafish chemical screening. Curr. Opin. Chem. Biol. 24, 58–70 (2015).

37. Arana, Á., Damiano, A. & Sánchez, L. Modeling neuropsychiatric diseases and drug responses in zebrafish. in Zebrafishno Model in Medical Research (ed. Disner, G. R.) (IntechOpen, London, 2025). doi:10.5772/intechopen.1011594.

38. Rosa, J. G. S., Lima, C. & Lopes-Ferreira, M. Zebrafish larvae behavior models as a tool for drug screenings and pre-clinical trials: a review. Int. J. Mol. Sci. 23, 6647 (2022).

39. Basnet, R. M., Zizioli, D., Taweedet, S., Finazzi, D. & Memo, M. Zebrafish larvae as a behavioral model in neuropharmacology. Biomedicines 7, 23 (2019).

40. Sampedro-Castañeda, M. et al. Epilepsy-linked kinase CDKL5 phosphorylates voltage-gated calcium channel Cav2.3, altering inactivation kinetics and neuronal excitability. Nat. Commun. 14, 7830 (2023).

41. Yan, M., Guo, X. & Xu, C. Revealing the complex role of CDKL5 in developmental epilepsy through a calcium channel related vision. Acta Epileptologica 6, 15 (2024).

42. Kim, E. K. & Choi, E. J. Pathological roles of MAPK signaling pathways in human diseases. Biochim. Biophys. Acta Mol. Basis Dis. 1802, 396–405 (2010).

43. Iroegbu, J. D., Ijomone, O. K., Femi-Akinlosotu, O. M. & Ijomone, O. M. ERK/MAPK signalling in the developing brain: Perturbations and consequences. Neurosci. Biobehav. Rev. 131, 792–805 (2021).

44. Kim, E. K. & Choi, E. J. Compromised MAPK signaling in human diseases: an update. Arch. Toxicol. 89, 867–882 (2015).

45. Fuchs, C. et al. Heterozygous CDKL5 knockout female mice are a valuable animal model for CDKL5 disorder. Neural Plast. 2018, 9726950 (2018).

46. Wang, I.-T. et al. Loss of CDKL5 disrupts kinome profile and event-related potentials leading to autistic-like phenotypes in mice. Proc Natl Acad Sci USA 109, 21516–21521 (2012).

47. Loi, M. et al. Increased DNA damage and apoptosis in CDKL5-deficient neurons. Mol. Neurobiol. 57, 2244–2262 (2020).

48. Zhu, Z. A. et al. CDKL5 deficiency in adult glutamatergic neurons alters synaptic activity and causes spontaneous seizures via TrkB signaling. Cell Rep. 42, 113202 (2023).

49. Trazzi, S. et al. HDAC4: A key factor underlying brain developmental alterations in CDKL5 disorder. Hum. Mol. Genet. 25, 3887–3907 (2016).

50. Khanam, T. et al. CDKL5 kinase controls transcription-coupled responses to DNA damage. EMBO J. 40, e108271 (2021).

51. Massey, S. et al. Novel CDKL5 targets identified in human iPSC-derived neurons. Cell. Mol. Life Sci. 81, 347 (2024).

52. Mari, F. et al. CDKL5 belongs to the same molecular pathway of MeCP2 and it is responsible for the early-onset seizure variant of Rett syndrome. Hum. Mol. Genet. 14, 1935–1946 (2005).

53. Kameshita, I. et al. Cyclin-dependent kinase-like 5 binds and phosphorylates DNA methyltransferase 1. Biochem. Biophys. Res. Commun. 377, 1162–1167 (2008).

54. Shukla, S. & Tekwani, B. L. Histone deacetylases inhibitors in neurodegenerative diseases, neuroprotection and neuronal differentiation. Front. Pharmacol. 11, 537 (2020).

55. Zhang, L. Y., Zhang, S. Y., Wen, R., Zhang, T. N. & Yang, N. Role of histone deacetylases and their inhibitors in neurological diseases. Pharmacol. Res. 208, 107410 (2024).

56. Kazantsev, A. G. & Thompson, L. M. Therapeutic application of histone deacetylase inhibitors for central nervous system disorders. Nat. Rev. Drug Discov. 7, 854–868 (2008).

57. Willmore, L. J., Shu, V. & Wallin, B. Efficacy and safety of add-on divalproex sodium in the treatment of complex partial seizures. Neurology 46, 49–53 (1996).

58. Beydoun, A., Sackellares, J. C. & Shu, V. Safety and efficacy of divalproex sodium monotherapy in partial epilepsy: A double-blind, concentration-response design clinical trial. Neurology 48, 182–188 (1997).

59. Watchon, M. et al. Sodium valproate increases activity of the sirtuin pathway resulting in beneficial effects for spinocerebellar ataxia-3 in vivo. Mol. Brain 14, 128 (2021).

60. Zádori, D., Geisz, A., Vámos, E., Vécsei, L. & Klivényi, P. Valproate ameliorates the survival and the motor performance in a transgenic mouse model of Huntington’s disease. Pharmacol. Biochem. Behav. 94, 148–153 (2009).

61. Yang, Q., Zhang, H., Jin, Z., Zhang, B. & Wang, Y. Effects of valproic acid therapy on rats with spinal cord injury: A systematic review and meta-analysis. World Neurosurg. 182, 12–28 (2024).

62. da Silva, H. C. et al. Anxiolytic and anticonvulsant effects of fisetin isolated from Bauhinia pentandra on adult zebrafish (Danio rerio). Chem. Biodivers. 21, e202401207 (2024).

63. Ahmad, A., Ali, T., Rehman, S. U. & Kim, M. O. Phytomedicine-based potent antioxidant, fisetin protects CNS-insult LPS-induced oxidative stress-mediated neurodegeneration and memory impairment. J. Clin. Med. 8, 850 (2019).

64. Flores, G., Vázquez-Roque, R. A. & Diaz, A. Resveratrol effects on neural connectivity during aging. Neural Regen. Res. 11, 1067–1068 (2016).

65. Almeida, E. R. et al. Micronized resveratrol shows anticonvulsant properties in pentylenetetrazole-Induced seizure model in adult zebrafish. Neurochem. Res. 46, 241–251 (2021).

66. Meng, X. J., Wang, F. & Li, C. K. Resveratrol is neuroprotective and improves cognition in pentylenetetrazole-kindling model of epilepsy in rats. Indian J. Pharm. Sci. 76, 125–131 (2014).

67. dos Santos, M. G., et al. Neuroprotective effects of resveratrol in in vivo and in vitro experimental models of Parkinson’s disease: a systematic review. Neurotox. Res. 40, 319–345 (2022).

68. Murray, K. O. et al. Intermittent supplementation with Fisetin improves physical function and decreases cellular senescence in skeletal muscle with aging: A comparison to genetic clearance of senescent cells and synthetic senolytic approaches. Aging Cell 24, e70114 (2025).

69. Liu, L. et al. Senolytic elimination of senescent macrophages restores muscle stem cell function in severely dystrophic muscle. Aging 14, 7650–7661 (2022).

70. Wang, T. H. et al. Fisetin exerts antioxidant and neuroprotective effects in multiple mutant hSOD1 models of amyotrophic lateral sclerosis by activating ERK. Neuroscience 379, 152–166 (2018).

71. Mancuso, R. et al. Resveratrol improves motoneuron function and extends survival in SOD1G93A ALS mice. Neurotherapeutics 11, 419–432 (2014).

72. Zhang, L. et al. Resveratrol alleviates motor and cognitive deficits and neuropathology in the A53T α-synuclein mouse model of Parkinson’s disease. Food Funct. 9, 6414–6426 (2018).

73. Kawamura, K. et al. Resveratrol improves motor function in patients with muscular dystrophies: an open-label, single-arm, phase IIa study. Sci. Rep. 10, 20585 (2020).

74. Guo, Y. J. et al. Resveratrol alleviates MPTP-induced motor impairments and pathological changes by autophagic degradation of α-synuclein via SIRT1-deacetylated LC3. Mol. Nutr. Food Res. 60, 2161–2175 (2016).

75. Ottappilakkil, H., Yesudas, G. H., Sreedharan, T. & Perumal, E. Fisetin modulates fluoride induced osteochondral toxicity in zebrafish larvae. Comp. Biochem. Physiol. C Toxicol. Pharmacol. 299, 110351 (2026).

76. Léotoing, L., Davicco, M. J., Lebecque, P., Wittrant, Y. & Coxam, V. The flavonoid fisetin promotes osteoblasts differentiation through Runx2 transcriptional activity. Mol. Nutr. Food Res. 58, 1239–1248 (2014).

77. Wang, X., Li, X., Zhou, J., Lei, Z. & Yang, X. Fisetin suppresses chondrocyte senescence and attenuates osteoarthritis progression by targeting sirtuin 6. Chem. Biol. Interact. 390, 110890 (2024).

78. Léotoing, L. et al. The polyphenol fisetin protects bone by repressing NF-κB and MKP-1-dependent signaling pathways in osteoclasts. PLoS One 8, e68388 (2013).

79. Han, X., Guo-Feng, J. & Zhu, F. Resveratrol alleviates osteoporosis by promoting osteogenic differentiation of bone marrow mesenchymal stem cells via SITR1/PI3K/AKT pathway. Int. J. Morphol 42, 216–224 (2024).

80. Zou, X., Xu, H. & Qian, W. The role and current research status of resveratrol in the treatment of osteoarthritis and its mechanisms: a narrative review. Drug Metab. Rev. 56, 399–412 (2024).

81. Zhao, W. et al. Effects of resveratrol on biochemical and structural outcomes in osteoarthritis: A systematic review and meta-analysis of preclinical studies. Heliyon 10, e34064 (2024).

82. Liao, W. & Lee, K. Z. CDKL5-mediated developmental tuning of neuronal excitability and concomitant regulation of transcriptome. Hum. Mol. Genet. 32, 3276–3298 (2023).

83. Van Bergen, N. J. et al. CDKL5 deficiency disorder: molecular insights and mechanisms of pathogenicity to fast-track therapeutic development. Biochem. Soc. Trans. 50, 1207–1224 (2022).

84. Panaretou, B. et al. Activation of the ATPase activity of Hsp90 by the stress-regulated cochaperone Aha1. Mol. Cell 10, 1307–1318 (2002).

85. Hussein, S. K., Bhat, R., Overduin, M. & LaPointe, P. Recruitment of Ahsa1 to Hsp90 is regulated by a conserved peptide that inhibits ATPase stimulation. EMBO Rep. 25, 3532–3546 (2024).

86. Chen, Y. et al. Type-I collagen produced by distinct fibroblast lineages reveals specific function during embryogenesis and Osteogenesis Imperfecta. Nat. Commun. 12, 7199 (2021).

87. Devos, H., Zoidakis, J., Roubelakis, M. G., Latosinska, A. & Vlahou, A. Reviewing the regulators of COL1A1. Int. J. Mol. Sci. 24, 10004 (2023).

88. Gistelinck, C. et al. Zebrafish collagen type I: molecular and biochemical characterization of the major structural protein in bone and skin. Sci. Rep. 6, 21540 (2016).

89. Raterman, S. T., Metz, J. R., Wagener, F. A. D. T. G. & Von den Hoff, J. W. Zebrafish models of craniofacial malformations: interactions of environmental factors. Front. Cell Dev. Biol. 8, 600926 (2020).

90. Ishikawa, D., Yamakita, S., Oh-Hashi, K. & Amaya, F. Critical role of p38α MAPK subclass in the development of pain hypersensitivity after hind paw incision. J. Pain Res. 18, 869–878 (2025).

91. O’Sullivan, S. A., Gasparini, F., Mir, A. K. & Dev, K. K. Fractalkine shedding is mediated by p38 and the ADAM10 protease under pro-inflammatory conditions in human astrocytes. J. Neuroinflammation 13, 189 (2016).

92. Chen, P. S. et al. Valproate protects dopaminergic neurons in midbrain neuron/glia cultures by stimulating the release of neurotrophic factors from astrocytes. Mol. Psychiatry 11, 1116–1125 (2006).

93. Fu, J. et al. Sodium valproate reduces neuronal apoptosis in acute pentylenetetrzole-induced seizures via inhibiting ER stress. Neurochem. Res. 44, 2517–2526 (2019).

94. Ueda, Y. & Willmore, L. J. Molecular regulation of glutamate and GABA transporter proteins by valproic acid in rat hippocampus during epileptogenesis. Exp. Brain Res. 133, 334–339 (2000).

95. Vriend, J. P. & Alexiuk, N. A. Effects of Valproate on amino acid and monoamine concentrations in striatum of audiogenic seizure-prone Balb/c mice. Mol. Chem. Neuropathol. 27, 307–324 (1996).

96. Loscher, W. & Honack, D. Comparison of anticonvulsant efficacy of valproate during prolonged treatment with one and three daily doses or continuous (’ “controlled release”) administration in a model of generalized seizures in rats. Epilepsia 36, 929–937 (1995).

97. Nissinen, J. & Pitkänen, A. Effect of antiepileptic drugs on spontaneous seizures in epileptic rats. Epilepsy Res. 73, 181–191 (2007).

98. Romoli, M. et al. Valproic acid and epilepsy: From molecular mechanisms to clinical evidences. Curr. Neuropharmacol. 17, 926–946 (2018).

99. Chen, J. et al. Developmental and behavioral alterations in zebrafish embryonically exposed to valproic acid (VPA): An aquatic model for autism. Neurotoxicol. Teratol. 66, 8–16 (2018).

100. Moore, S. J. et al. A clinical study of 57 children with fetal anticonvulsant syndromes. J Med Genet 37, 489–497 (2000).

101. Wagner, G. C., Reuhl, K. R., Cheh, M., McRae, P. & Halladay, A. K. A new neurobehavioral model of autism in mice: Pre- and postnatal exposure to sodium valproate. J. Autism Dev. Disord. 36, 779–793 (2006).

102. Schneider, T. & Przewłocki, R. Behavioral alterations in rats prenatally to valproic acid: Animal model of autism. Neuropsychopharmacology 30, 80–89 (2005).

103. Williams, G. et al. Fetal valproate syndrome and autism: additional evidence of an association. Dev. Med. Child Neurol. 43, 202–206 (2001).

104. Khatoon, S., Samim, M., Dahalia, M. & Nidhi. Fisetin provides neuroprotection in pentylenetetrazole-induced cognition impairment by upregulating CREB/BDNF. Eur. J. Pharmacol. 944, 175583 (2023).

105. Chuang, J. Y. et al. Regulatory effects of fisetin on microglial activation. Molecules 19, 8820–8839 (2014).

106. Stockinger, J., Maxwell, N., Shapiro, D., DeCabo, R. & Valdez, G. Caloric restriction mimetics slow aging of neuromuscular synapses and muscle fibers. Journals of Gerontology - Series A Biological Sciences and Medical Sciences 73, 21–28 (2018).

107. Al Sagheer, T. et al. Motor impairments correlate with social deficits and restricted neuronal loss in an environmental model of autism. International Journal of Neuropsychopharmacology 21, 871–882 (2018).

108. Bath, K. G. & Pimentel, T. Effect of early postnatal exposure to valproate on neurobehavioral development and regional BDNF expression in two strains of mice. Epilepsy and Behavior 70, 110–117 (2017).

109. Shakibaei, M., Buhrmann, C. & Mobasheri, A. Resveratrol-mediated SIRT-1 interactions with p300 modulate receptor activator of NF-κB ligand (RANKL) activation of NF-κB Signaling and inhibit osteoclastogenesis in bone-derived cells. Journal of Biological Chemistry 286, 11492–11505 (2011).

110. Ahmad Hairi, H., Jayusman, P. A. & Shuid, A. N. Revisiting resveratrol as an osteoprotective agent: Molecular evidence from in vivo and in vitro studies. Biomedicines 11, 1453 (2023).

111. Feng, J. et al. Protective effects of resveratrol on Postmenopausal osteoporosis: Regulation of SIRT1-NF-κB signaling pathway. Acta Biochim. Biophys. Sin. (Shanghai). 46, 1024–1033 (2014).

112. Damjanov, N., Kauffman, R. S. & Spencer-Green, G. T. Efficacy, pharmacodynamics, and safety of VX-702, a novel p38 MAPK inhibitor, in rheumatoid arthritis: Results of two randomized, double-blind, placebo-controlled clinical studies. Arthritis Rheum. 60, 1232–1241 (2009).

113. Ding, C. Drug evaluation: VX-702, a MAP kinase inhibitor for rheumatoid arthritis and acute coronary syndrome. Current Opinion in Investigational Drugs 7, 1020–1025 (2006).

114. Boluk, A. et al. The effect of valproate on bone mineral density in adult epileptic patients. Pharmacol. Res. 50, 93–97 (2004).

115. Fan, D., Miao, J., Fan, X., Wang, Q. & Sun, M. Effects of valproic acid on bone mineral density and bone metabolism: A meta-analysis. Seizure 73, 56–63 (2019).

116. Humphrey, E. L., Morris, G. E. & Fuller, H. R. Valproate reduces collagen and osteonectin in cultured bone cells. Epilepsy Res. 106, 446–450 (2013).

117. Xie, X. et al. Bone-targeting engineered small extracellular vesicles carrying anti-miR-6359-CGGGAGC prevent valproic acid-induced bone loss. Signal Transduct. Target. Ther. 9, 24 (2024).

118. Fan, H. C., Wang, S. Y., Peng, Y. J. & Lee, H. S. Valproic acid impacts the growth of growth plate chondrocytes. Int. J. Environ. Res. Public Health 17, 3675 (2020).

119. Paradis, F. H. & Hales, B. F. Exposure to valproic acid inhibits chondrogenesis and osteogenesis in mid-organogenesis mouse limbs. Toxicological Sciences 131, 234–241 (2013).

120. Gebuijs, I. G. E. et al. The anti-epileptic drug valproic acid causes malformations in the developing craniofacial skeleton of zebrafish larvae. Mech. Dev. 163, 103632 (2020).

121. Bradley, E. W., Mcgee-Lawrence, M. E., Westendorf, J. J., Clinic, M. & Rochester, M. Hdac-mediated control of endochondral and intramembranous ossification. Critical Reviews TM in Eukaryotic Gene Expression 21, 101–113 (2011).

