## Supplementary Table S1 for "Rescuing functional defects in a zebrafish model of CDKL5 deficiency disorder: Contribution to the identification of new therapeutic compounds"

**Supplementary Table S1**. List of 170 compounds screened for their potential to rescue locomotor behavior in *cdkl5*$^{-/-}$ zebrafish.

| MAPK Inhibitor library | | | |
|---|---|---|---|
| 1 | Selumetinib | 44 | AZD8330 |
| 2 | SB202190 | 45 | Apigenin |
| 3 | Ginkgolide B | 46 | TAK-733 |
| 4 | BIX 02189 | 47 | WHI-P154 |
| 5 | SB590885 | 48 | Quercitrin |
| 6 | Honokiol | 49 | TBHQ |
| 7 | PH-797804 | 50 | SL-327 |
| 8 | Notoginsenosi | 51 | PD98059 |
| 9 | Lidocaine hydrochloride | 52 | Pimasertib |
| 10 | Pseudolaric Acid B | 53 | RAF265 |
| 11 | BMS-S36924 | 54 | Asiatic Acid |
| 12 | Refametinib | 55 | A-674563 |
| 13 | Lidocaine | 56 | GW5074 |
| 14 | AT7867 | 57 | Baohuoside I |
| 15 | Palomid 529 | 58 | Dabrafenib Mesylate |
| 16 | Myricetin | 59 | WHI-P258 |
| 17 | AZ 628 | 60 | SB431542 |
| 18 | Guggulsterone E&Z | 61 | Regorafenib |
| 19 | Trans-Zeatin | 62 | Hesperadin |
| 20 | Ferulic acid methyl ester | 63 | PF-4708671 |
| 21 | PD184352 | 64 | Glycyrrhizin |
| 22 | U0126-EtOH | 65 | Trametinib |
| 23 | Resveratrol | 66 | TAK-715 |
| 24 | PD318088 | 67 | Isopsoralen |
| 25 | (-)-Epigallocatechin Gallate | 68 | Regorafenib Monohydrate |
| 26 | Myricitrin | 69 | B-Raf IN 1 |
| 27 | Dabrafnib | 70 | SB203580 |
| 28 | Ginsenoside Re | 71 | Vemurafenib |
| 29 | JNK-IN-8 | 72 | BIX 02188 |
| 30 | VX-702 | 73 | Raf265 derivative |
| 31 | PD0325901 | 74 | Hesperetin |
| 32 | GDC-0879 | 75 | ZM 336372 |
| 33 | VX-745 | 76 | Methylthiouracil |
| 34 | Doramapimod | 77 | Astragaloside IV |
| 35 | Esculin | 78 | Osimertinib mesylate |
| 36 | Bupivacaine HCL | 79 | Longdaysin |
| 37 | BI-D1870 | 80 | Binimetinib (MEK162) |
| 38 | Bakuchiol | 81 | Osimertinib (AZD9291) |
| 39 | Skatole | 82 | XDM8-92 |
| 40 | PF-6260933 | 83 | VX-11e |
| 41 | Sorafenib Tosylate | 84 | LJH685 |
| 42 | PLX-4720 | 85 | CEP-32496 |
| 43 | SP600125 | 86 | eFT-508 |

| | | | |
|---|---|---|---|
| 87 | Cu-CPT22 | 119 | Beta-Elemonic |
| 88 | Hydroxy safflor yellow A | 120 | Bisindolylmaleimide IX |
| 89 | Shanzhiside methyl ester | 121 | Anisomycin |
| 90 | MLN2480 | 122 | DTP3 |
| 91 | ERK5-IN-1 | 123 | Pexmetinib |
| 92 | Piperlongumine | 124 | SIS3 HCL |
| 93 | SB239063 | 125 | BAW2881 |
| 94 | LJI308 | 126 | Tanzisrtib |
| 95 | Cobimetinib | 127 | LXH254 |
| 96 | NQDI-1 | 128 | Polyphyllin I |
| 97 | RAF709 | 129 | Dehydrocorydalin |
| 98 | Anhydroicaritin | 130 | Skepinone-L |
| 99 | 2',5' Dihydroxyacetophenone | 131 | CGP 57380 |
| 100 | TIC10 Analogue | 132 | OTS964 |
| 101 | URMC-099 | 133 | Licochalcone A |
| 102 | GDC-0623 | 134 | TIC10 |
| 103 | CCT196969 | 135 | ML264 |
| 104 | Sodium Tauroursodeoxycholate | 136 | AD80 |
| 105 | BMS-582949 | 137 | AZ304 |
| 106 | Selonsertib | 138 | Carnosol |
| 107 | MK-8353 | 139 | Isorhamnetin 3-O-neohesperoside |
| 108 | 10-Gingerol | 140 | Losmapimod |
| 109 | Lingustilide | 141 | JNK Inhibitor IX |
| 110 | RO5126766 | 142 | OTS514 hydrochloride |
| 111 | Sorafenib | 143 | BI-847325 |
| 112 | AT13148 | 144 | PLX7904 |
| 113 | Pluripotin | 145 | BI-78D3 |
| 114 | DEL-22379 | 146 | LY3214996 |
| 115 | Pamapimod | 147 | CC-90003 |
| 116 | APS-2-79 HCL | 148 | Nitidine Chloride |
| 117 | UM-164 | 149 | Fraxetin |
| 118 | Norisoboldine | | |

**Histone Modification library**

| | | | |
|---|---|---|---|
| 1 | Panobinostat | 12 | ZM 447439 |
| 2 | MLN8054 | 13 | SNS-314 |
| 3 | Barasertib | 14 | Droxinostat |
| 4 | Resveratrol | 15 | Curcumin |
| 5 | Divalproex sodium | 16 | (-)-Parthenolide |
| 6 | Fisetin | 17 | GSK1070916 |
| 7 | AMG-900 | 18 | Entacapone |
| 8 | Bufexamac | 19 | Sodium Phenylbutyrate |
| 9 | Tolcapone | 20 | Amodiaquine hydrochloride |
| 10 | AMI-1, free acid | 21 | Tozasertib |
| 11 | Vorinostat | | |
